# Genetic variants within silencer elements shape human blood cell phenotype diversity

**DOI:** 10.1101/2025.01.13.632723

**Authors:** Minkang Tan, Henk van Weerd, Alper Yavas, Baoxu Pang

## Abstract

Non-coding variants contribute to blood cell trait variation, but their effects on silencers remain poorly defined. Here, we used ReSE, a high-throughput silencer screening system, to assess sequences encompassing 14,720 non-coding variants from fine-mapped loci associated with 15 blood cell traits. In K562 cells, we identified 805 variants with differential silencer activity, comprising 425 loss-of-silencer-function and 380 gain-of-silencer-function candidates. Luciferase reporter assays confirmed allele-dependent repression, and endogenous editing of selected variants was accompanied by altered expression of nearby genes. K562 cells harboring the rs2857078 alternative allele showed downregulated globin expression and reduced mature erythrocyte markers upon erythroid differentiation. By integrating single-cell hematopoietic chromatin accessibility with evolutionary conservation, we further prioritized functional silencer variants despite their low fine-mapping posterior inclusion probabilities. Together, these findings implicate silencer perturbation as a regulatory mechanism contributing to blood cell trait variation and related disorders, and provide candidates for mechanistic investigation of non-coding associations.

## Introduction

Non-coding regulatory elements (NCRE) control the timing, magnitude and cellular specificity of gene expression^1^. Although promoters, enhancers and insulators have been extensively studied^2–8^, silencers remain less comprehensively characterized. Silencers are cis-regulatory elements that repress transcription, often through the recruitment of transcriptional repressors and corepressors^9–12^. Accessible chromatin can harbor silencers^9,10,13,14^, but accessibility alone does not distinguish repressive from activating regulatory activity^14,15^. Recent computational and functional screens have expanded silencer catalogs and revealed diverse chromatin features and mechanisms of repression^9–17^. Nevertheless, how genetic variation alters silencer activity and contributes to human trait variation remains incompletely understood.

Many genetic variants associated with complex traits and diseases lie outside protein-coding sequences, making their functional interpretation challenging^18–21^. Non-coding regulatory variants can alter transcription factor binding sites and thereby influence gene expression^19,21^. Large-scale functional studies have identified allele-dependent regulatory activity at numerous GWAS loci, particularly in enhancers^22–25^. By comparison, the effects of genetic variation on silencer activity remain less well characterized. Variants that weaken silencer activity could instead increase target gene expression, whereas variants that strengthen or create repressive activity could reduce target-gene expression. Systematically evaluating these effects would extend functional interpretation of GWAS loci beyond enhancer-mediated regulation. However, incomplete silencer annotation and limited characterization of allelic silencer activity have constrained this effort.

Blood cell traits provide a well-characterized setting to investigate the effects of genetic variation on NCRE function, including silencers. Blood cell counts and indices reflect aspects of hematopoietic differentiation, cell production and maintenance, and hemoglobin biology^26,27^. Extensive genetic and epigenomic resources also enable regulatory variants to be investigated across hematopoietic cell populations^28–31^. The Blood Cell Consortium characterized 15 blood cell traits in 746,667 individuals from five ancestry groups and identified 5,552 trait–variant associations^28^. These traits encompass erythrocyte, leukocyte and platelet counts, leukocyte subtype counts, and measures of erythrocyte size, hemoglobin content and platelet volume. Together, these resources provide a foundation for testing potential silencer activity at blood-trait-associated loci and investigating candidate target genes and relevant cellular contexts.

Here we used ReSE^9^ to screen both alleles of 14,720 non-coding variants from fine-mapped loci associated with 15 blood cell traits in K562 and HepG2 cells. We identified candidates associated with loss or gain of silencer function and used motif analysis to predict changes in transcription factor binding sites. We integrated these findings with whole-blood cis-eQTL data^29^, hematopoietic chromatin accessibility profiles^31^ and promoter-capture Hi-C maps^30^ to nominate candidate target genes and cellular contexts. Reporter assays supported allele-dependent repression, and endogenous genome editing of selected variants was accompanied by altered local gene expression. Finally, we integrated ChromBPNet-derived predictions of hematopoietic regulatory effects^32^ with evolutionary conservation using a FLARE-based framework^33^ to prioritize candidate functional silencer variants. These analyses identify allelic silencer activity as a mechanism to consider when interpreting non-coding blood-trait associations.

## Results

### Selection of blood-trait-associated variants for silencer screening

Functional studies of non-coding variation have provided insights into the regulation of blood cell traits and hematological diseases^34–36^. For example, dissection of the BCL11A erythroid enhancer has identified regulatory sequences with therapeutic potential for disorders of hemoglobin production^37^. Defining the functional effects of blood-trait-associated variants can therefore help connect genetic associations to regulatory mechanisms. Compared with enhancers, whose annotation is aided by features such as H3K27ac enrichment and occupancy by the transcriptional coactivator p300^38^, silencers remain less comprehensively characterized. We therefore sought to assess the effects of non-coding GWAS variants on silencer activity using a functional screen. Such variants could weaken repression by disrupting repressor-binding sites or strengthen repression by creating or enhancing such sites (**Fig. 1a**).

**Figure 1.**
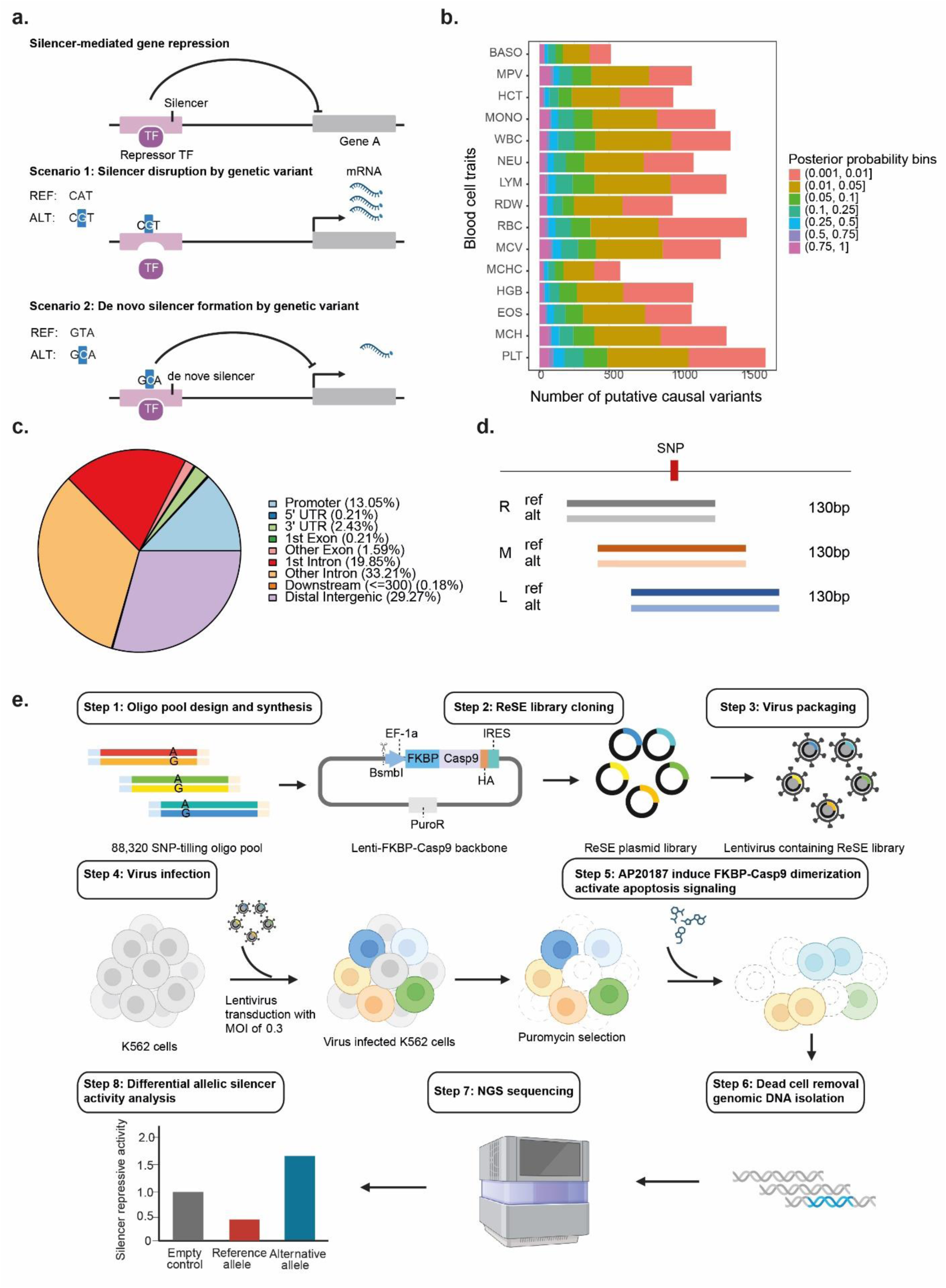
Prioritization of candidate non-coding GWAS variants affecting silencer element function. **(a).** Schematic representation of genetic variants such as SNPs can disrupt or create silencer elements to alter target gene repression. Upper panel: silencer-mediated gene repression. The silencer element recruits a repressor TF, which suppresses the transcription of target gene. Middle panel: silencer disruption. A SNP within the silencer disrupts the repressor TF binding motif, abolishing TF binding and resulting in de-repression of the target gene. Lower panel: de novo silencer formation. A SNP creates a new repressor TF binding motif, generating a de novo silencer element that represses target gene expression. **(b).** Prioritization of blood trait-associated SNPs for silencer function evaluation. SNPs were prioritized by fine-mapping within 95% credible sets across 15 blood cell traits from the trans-ancestry meta-analysis. Bars are stacked by bins of each variant’s highest posterior probability of being causal. **(c).** Genomic annotation of blood trait-associated SNPs for silencer function evaluation. All included variants were categorized by genomic feature using Ensembl Variant Effect Predictor (VEP) annotation. **(d).** Oligonucleotide design for tiling GWAS variants in the ReSE silencer screen assay. For each SNP, a pair of 130-nt oligonucleotides was designed, carrying either the reference (ref) or alternate (alt) allele at the variant site. Variants were tiled at three positions along the oligonucleotide: one-quarter (left), one-half (middle), and three-quarters (right) of the total length, to control for position effects within the oligonucleotide. **(e)** Overview of the ReSE SNP screen. An oligonucleotide array was designed and synthesized to carry both allelic bases of each SNP, with the SNP tiled at different positions along the oligonucleotide. The PCR-amplified oligonucleotide library was digested with BsmBI and cloned into the lentiviral screening vector pLenti-FKBP-delCasp9-Puro. The resulting ReSE library was transfected into 293T cells together with packaging plasmids to produce the lentiviral SNP ReSE library. K562 cells were transduced at the MOI of 0.3 and selected with puromycin. Upon addition of a dimerizer, expressed caspase 9 is activated and induces apoptosis; cells survive only when the inserted SNP-containing fragment acts as a silencer and represses caspase 9 expression. Fragments enriched in surviving cells (post-dimerizer) relative to untreated cells (pre-dimerizer) were identified as candidate silencers, and statistical analysis was used to quantify differential allelic silencer activity attributable to each SNP.

We selected 14,720 non-coding variants from fine-mapped loci associated with at least one of the 15 blood cell traits (**Fig. 1b**), drawing on a genetic association study spanning European, South Asian, Hispanic, East Asian and African ancestry groups^28^. Following annotation-based filtering using the Ensembl Variant Effect Predictor (VEP), most selected variants were located in intronic or intergenic regions (**Fig. 1c**). Blood cell traits provide a useful setting for this analysis because of their polygenic architecture, relevance to human disease and extensive genetic and regulatory characterization. We used K562, a human chronic myeloid leukemia cell line previously used to investigate erythroid regulatory variation^25^, as the cellular model for the initial screen. This system enabled us to test allele-dependent repression in a hematopoietic cellular context.

We adapted the ReSE silencer screening system^9^ to assay 130-bp DNA fragments encompassing each selected variant. For each variant, we designed three overlapping 130-bp fragments with the variant positioned approximately one-quarter, one-half or three-quarters along the fragment, corresponding to the “left,” “middle” and “right” windows, respectively **(Fig. 1d**). Both reference and alternative alleles were synthesized for each window, yielding six constructs per variant and 88,320 variant-containing fragments. This design sampled different flanking sequences to increase the opportunity to capture sequence features contributing to repression. We supplemented these fragments with a reported 3,446 silencer control sequences selected from SilencerDB^39^ and cloned the resulting pooled library, termed ReSE_SNP, into the ReSE screening vector. The reported final library comprised 91,766 DNA fragments.

### High-throughput identification of silencer activity at blood-trait-associated loci

We performed ReSE screening as previously described^9^. The ReSE_SNP library was transduced into K562 cells in three biological replicates, and genomic DNA was collected from the initial cell populations before selection (Ctrl) and from surviving cells after AP20187-induced apoptosis, which should enrich for potential silencer fragments (AP; **Fig. 1e**). Library inserts were amplified from genomic DNA and quantified by next-generation sequencing. Most library fragments remained detectable in the screened populations (**Fig. 2a**), and fragment abundance was correlated across biological replicates (**Fig. 2b**). We assessed enrichment after selection using MAGeCK, which models construct counts using a negative binomial distribution^40^. This analysis considered constructs carrying either reference or alternative alleles and identified 9,629 candidate variant-associated silencers at a false discovery rate (FDR) below 0.01.

**Figure 2.**
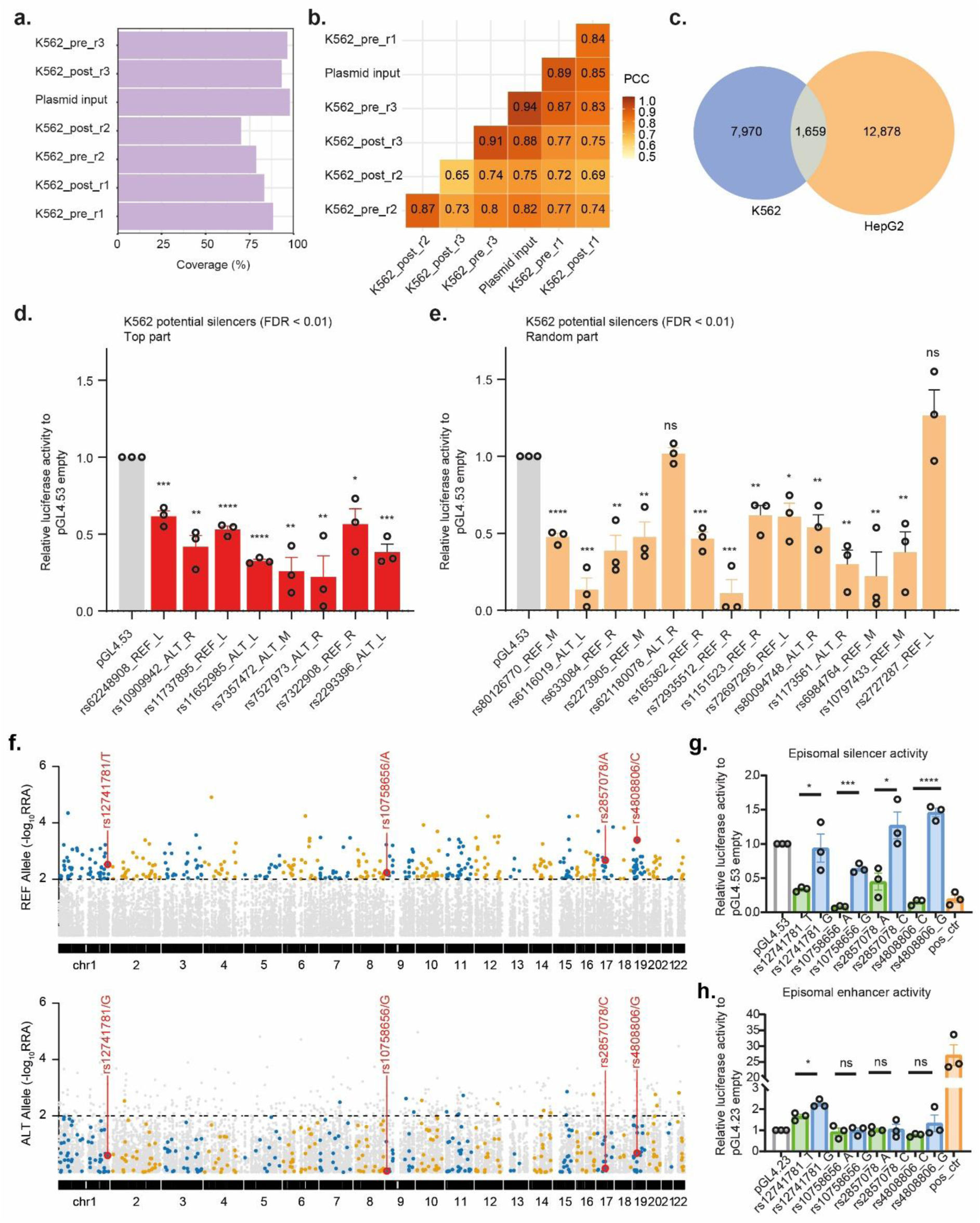
Identification of silencer elements residing in GWAS variants loci. **(a)** Coverage of ReSE SNP library complexity in the plasmid library input and in 3 biological replicates of screening samples (both pre- and post-dimerizer treatment) in K562 cells. **(b)** Pearson correlation between biological replicates of the ReSE SNP libraries isolated from K562 cells (pre: cell population before dimerizer treatment; post: surviving cells after dimerizer addition; input: constructed ReSE plasmid library). **(c)** Venn diagram of SNP-containing silencers identified in K562 and HepG2 cells using MAGeCK-RRA at an FDR threshold of < 0.01. **(d, e)** Luciferase assays validating the repressive activity of SNP-containing silencers identified in K562 cells (FDR < 0.01). Colors indicate silencers selected from the top (d) or randomly from (**e**) the FDR-ranked list. Silencer regions were PCR-amplified from the input plasmid library DNA and cloned upstream of the promoter in the firefly luciferase reporter plasmid pGL4.53; the empty pGL4.53 plasmid served as the baseline control. pGL4.53 and Renilla plasmids were co-transfected into HepG2 cells by electroporation. The y axis shows luciferase activity as a percentage of the empty pGL4.53 control (n = 3 biologically independent samples; bars show mean ± s.e.m.; ns, P > 0.05; *P < 0.05; **P < 0.01; ***P < 0.001; ****P < 0.0001; two-tailed unpaired t-test). **(f)** Rank difference test statistical framework to identify GWAS SNPs with differential allelic silencer activity. The combined Manhattan plot shows the distribution of all SNPs included in the ReSE screen, with reference (upper panel) and alternate (lower panel) allelic silencer activity shown. Significant hits fall above the dashed line, corresponding to a MAGeCK-RRA score cutoff of 0.01. SNPs selected for downstream analysis are indicated. **(g)** Luciferase assays to determine the effect of GWAS variants on silencer activity. SNP-containing fragments from the ReSE screen were PCR-amplified and cloned into the silencer reporter plasmid pGL4.53, driven by a PGK promoter. The empty pGL4.53 plasmid served as the baseline control. The y axis shows luciferase activity in relative units compared with the empty pGL4.53 plasmid (n = 3 biologically independent samples; bars show mean ± s.d.; *P < 0.05, ***P < 0.001, ****P < 0.0001; two-tailed unpaired t-test). **(h)** Luciferase assays to determine the effect of GWAS variants on enhancer activity. SNP-containing fragments from the ReSE screen were PCR-amplified and cloned into the enhancer reporter plasmid pGL4.23, driven by a minimal promoter. The empty pGL4.23 plasmid served as the baseline control. The y axis shows luciferase activity in relative units compared with the empty pGL4.23 plasmid (n = 3 biologically independent samples; bars show mean ± s.d.; ns, P > 0.05; *P < 0.05; two-tailed unpaired t-test).

To investigate whether silencer activity at blood-trait-associated loci differed between cellular contexts, we screened the same ReSE_SNP library in HepG2 cells, a well-characterized human hepatoblastoma cell line. Screening was performed in three biological replicates, with broad library representation and correlated fragment counts across replicates (**Supplementary Fig. 1a,b**). HepG2 provided a complementary hepatic context for investigating these sequences, motivated in part by the contribution of hepatic iron regulation^41^ and thrombopoietin production^42^ to blood cell homeostasis. The screen identified 14,537 candidate variant-associated silencers at FDR < 0.01. Of these, 1,659 were also detected in both K562 and HepG2 cells, representing 17.2% of HepG2 candidates and 11.4% of K562 candidates (**Fig. 2c**). Thus, the screens identified a shared subset of candidates, alongside larger sets detected in only one of the two cell lines.

To evaluate the repressive activity of candidate silencers selected at FDR < 0.01, we performed luciferase reporter assays on 8 top-ranked and 14 randomly selected candidate silencers from the K562 screen. All 8 top-ranked candidates and 12 of the 14 randomly selected candidates significantly reduced luciferase activity relative to the empty-vector control (**Fig. 2d,e**). We also tested 13 candidates from the HepG2 screen, comprising four top-ranked candidates, four candidates near the significance cutoff and five randomly selected candidates. All 13 significantly reduced luciferase activity relative to the empty-vector control (**Supplementary Fig. 1c**). These experiments supported the repressive activity of selected candidates identified in both screens. Together, the screening and validation results identify blood-trait-associated sequences with silencer activity and reveal differences in candidate detection between the two cellular contexts.

### Blood-trait-associated variants exhibit allele-specific silencer activity

To identify blood-trait-associated variants with differential silencer activity, we developed a rank-based framework that integrated evidence across the three tiled sequence windows (**Methods**). For each window, we calculated the difference between the ranks of the reference-and alternative-allele constructs obtained from the initial MAGeCK analysis^40^. These rank differences were then ranked and converted to percentiles. We applied a modified α-robust rank aggregation (α-RRA) procedure to combine the window-level evidence and prioritize variants with differential allelic activity (**Supplementary Fig. 2**). Statistical significance was assessed by permutation testing, followed by correction for multiple testing (**Methods**). We identified 425 loss-of-silencer-function (LoSF) and 380 gain-of-silencer-function (GoSF) candidates in K562 cells (**Fig. 2f**), and 387 LoSF and 409 GoSF candidates in HepG2 cells.

Four LoSF variants from K562 ReSE screen, rs12741781, rs10758656, rs2857078 and rs4808806, which are associated with red blood cell or platelet traits, were further tested by luciferase reporter assay (**Fig. 2g**). For each variant, fragments carrying the reference or alternative allele were cloned into the silencer luciferase reporter containing a PGK promoter. All four alternative-allele constructs produced significantly higher luciferase activity than their corresponding reference-allele constructs, consistent with reduced repression and supporting the direction of the effects identified by ReSE (**Fig. 2g**). To investigate whether increased reporter expression could also reflect allele-dependent enhancer activity, we tested the same variant-containing sequences in a seperate enhancer reporter containing a minimal promoter. Three candidates, rs10758656, rs2857078 and rs4808806, showed no significant difference between alleles in this assay. By contrast, rs12741781 showed a modest but significant increase in reporter activity for the alternative allele (**Fig. 2h**). Thus, the reporter assays supported reduced silencer activity by the alternative allele at all four loci, while also indicating that rs12741781 can exhibit allele-dependent activating activity in a different promoter context.

### Regulatory features and heritability enrichment of silencer-altering variants

We hypothesized that some allelic differences in repression could arise through weakening or strengthening transcription factor binding sites (**Fig. 3a**). Using motifbreakR^43^, we predicted allele-dependent changes in motif scores and examined recurrently affected transcription factor motifs. In K562 cells, LoSF variants were associated with lower alternative-allele motif scores for NF-κB family members, including NFKB1, REL and RELA. These factors participate in immune-cell signaling and hematopoietic regulation^44^. GoSF variants included predicted increases in motif scores for GFI1B, an essential regulator of erythroid and megakaryocytic development^45,46^. In HepG2 cells, LoSF variants showed predicted decreases in motif scores for HNF1A, RFX5 and IRF-family members, whereas GoSF variants showed predicted increases in motif scores for NR1D1, ZBTB7A and CEBPB (**Supplementary Fig. 3a**). These predictions nominate transcription factors that involve in blood cell traits regulation by affecting allele-dependent silencer regulatory activities.

**Figure 3.**
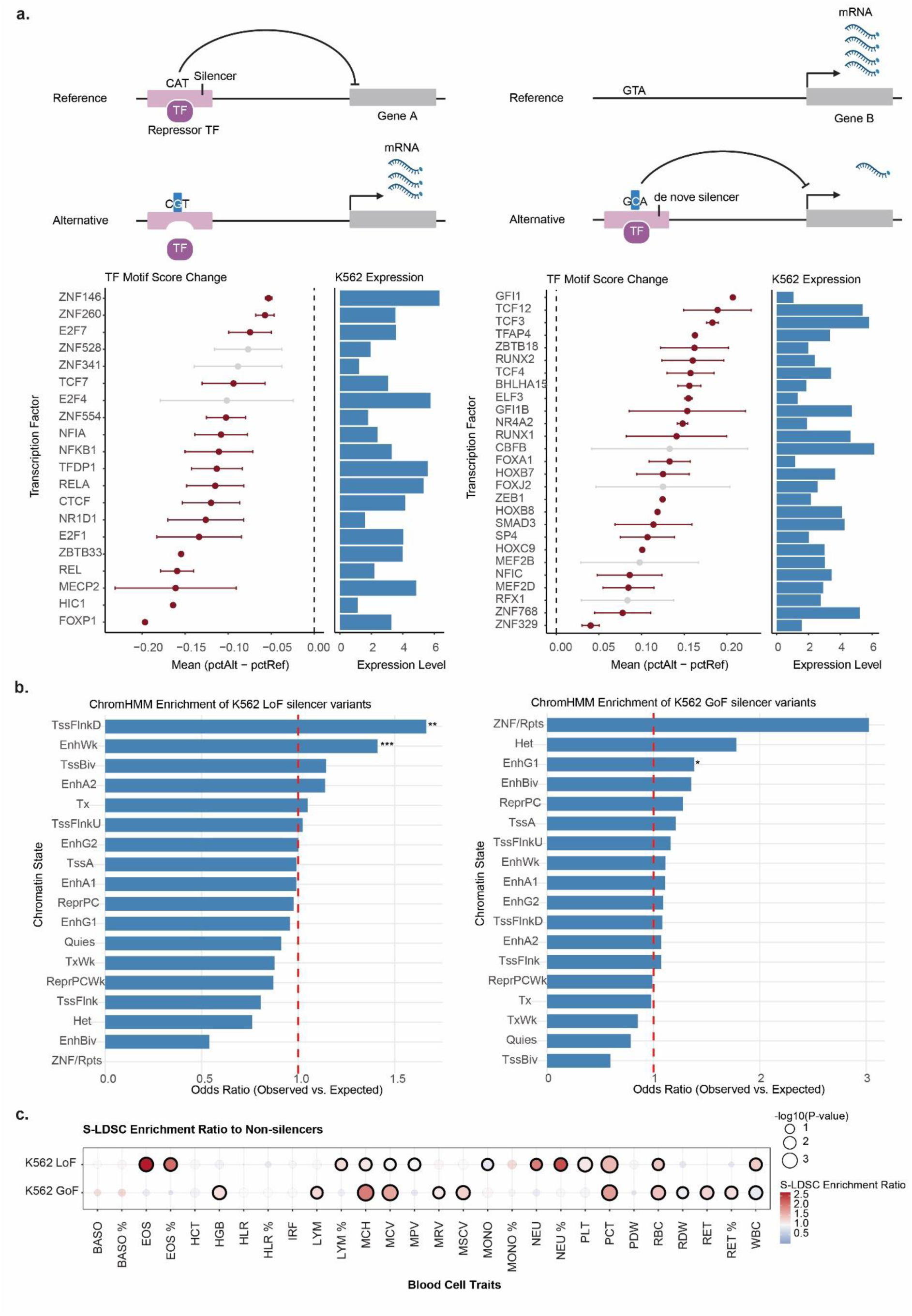
Epigenomic patterns and partitioned heritability of silencer-affecting GWAS variants. **(a)** Disruptive TF binding sites associated with LoSF and GoSF variants. Upper panel: schematic of how GWAS variants disrupt TF motifs, leading to LoF or GoF of silencer elements. Lower panel: recurrent disruptive TF motifs among LoSF and GoSF variants, scored and ranked with motifbreakR. The x axis shows the difference between the pctAlt and pctRef scores, the percentage of the best score each motif could achieve on an ideal reference or alternate sequence. TF motifs on the y axis are ranked by the mean difference between pctAlt and pctRef scores; error bars represent 95% confidence intervals. Significant transcription factor binding disruptions are highlighted in dark maroon, and non-significant factors are shown in light gray. RNA-seq expression levels of the TF genes in K562 cells, normalized as log(TPM + 1), are shown on the right. **(b)** Chromatin states associated with LoSF and GoSF variants. Enrichment of ChromHMM states within LoSF or GoSF variants was calculated relative to a background of all SNPs included in the ReSE screen. The red dashed line indicates the odds-ratio cutoff, comparing the observed frequency of each of the 18 ChromHMM states among LoSF or GoSF variants with the expected frequency. P values were calculated using Fisher’s exact test; *P < 0.05, **P < 0.01, ***P < 0.001. Chromatin state abbreviations: TssA, active transcription start site; TssFlnk, flanking TSS; TssFlnkU, flanking TSS upstream; TssFlnkD, flanking TSS downstream; Tx, strong transcription; TxWk, weak transcription; EnhA1/EnhA2, active enhancer; EnhG1/EnhG2, genic enhancer; EnhWk, weak enhancer; TssBiv, bivalent TSS; EnhBiv, bivalent enhancer; ReprPC, repressed Polycomb; ReprPCWk, weakly repressed Polycomb; Quies, quiescent/low; Het, heterochromatin; ZNF/Rpts, ZNF genes and repeats. **(c)** Enrichment of blood cell trait heritability in functional silencer variants. Heritability enrichment for blood cell traits within LoSF and GoSF variants was estimated by S-LDSC, which quantifies the heritability (h²) enrichment of SNPs in each annotation (LoSF, GoSF, and non-silencer variants). Relative enrichment of LoSF or GoSF variants is shown as a ratio to non-silencer variants. Bubble size indicates −log_10_(P); highlighted bubbles indicate significant enrichment (P < 0.05).

We next examined the chromatin environments of LoSF and GoSF variants. Chromatin-state annotations integrate histone modification patterns to distinguish promoter-associated, enhancer-associated, transcribed and repressed regions^47–49^. Using an 18-state segmentation, we compared the representation of these states among LoSF and GoSF variants with that among all screened variants. In K562 cells, LoSF variants showed enrichment in downstream transcription-start-site flanking regions (TssFlnkD) and weak enhancer regions (EnhWk), whereas GoSF variants showed enrichment in the genic enhancer state EnhG1 (**Fig. 3b**). The HepG2 analysis showed a different pattern, with enrichment in the weak Polycomb-repressed state ReprPCWk among LoSF variants and in EnhWk among GoSF variants at the nominal significance threshold shown in (**Supplementary Fig. 3b**). Thus, variants associated with differential silencer activity occurred in several chromatin environments, including regions annotated as enhancers.

Blood cell traits are highly polygenic, and functional genomic annotations can account for disproportionate shares of SNP heritability^50–52^. To examine the relationship between silencer-associated variants and blood-trait genetics, we used stratified LD score regression to estimate heritability enrichment for annotations defined by the K562 LoSF, GoSF and non-silencer variants (**Fig. 3c and Supplementary Fig. 3c**). Relative to the non-silencer annotation, LoSF variants showed higher enrichment estimates for eosinophil and neutrophil counts and percentages, whereas GoSF variants showed higher estimates for reticulocyte count and percentage. Both classes showed higher estimates for mean corpuscular hemoglobin, plateletcrit and red blood cell count. These patterns suggest that the two classes capture partly distinct, but overlapping, components of blood-trait-associated regulatory variation that related to silencer activity.

The observed trait patterns are broadly compatible with the hematopoietic functions of several transcription factors implicated by the motif analysis, including NF-κB family members^53,54^, RUNX1^55^ and GFI1B^46^. Together, these analyses identify diverse regulatory contexts for silencer-altering variants and suggest that variants with opposing effects on repression may contribute to shared and distinct aspects of blood cell trait variation.

### Linking silencer-altering variants to cellular contexts and target genes

A central challenge in interpreting non-coding GWAS variants is identifying the endogenous genes they regulate and the cellular contexts in which they act. We first investigated *rs2857078*, a candidate causal variant for mean corpuscular hemoglobin (MCH; fine-mapping posterior probability, PP = 0.398), red blood cell distribution width (RDW; PP = 0.788) and mean corpuscular hemoglobin concentration (MCHC; PP = 0.03) (**Fig. 4a**). This variant lies within an intron of SLC4A1, in a region showing erythroid-specific chromatin accessibility and overlap with H3K27me3 enrichment (**Supplementary Fig. 4a**). Hi-C data identified contacts between this region and the promoters of SLC25A39 and RUNDC3A in erythroblasts (**Fig. 4a**). MotifbreakR further predicted that *rs2857078* disrupts the binding site for NR1D1 (**Fig. 4b**), a constitutive transcriptional repressor that silences target gene expression by recruiting the nuclear receptor corepressor (NCoR)–HDAC3 complex, leading to histone deacetylation and chromatin compaction^56–58^.

**Figure 4.**
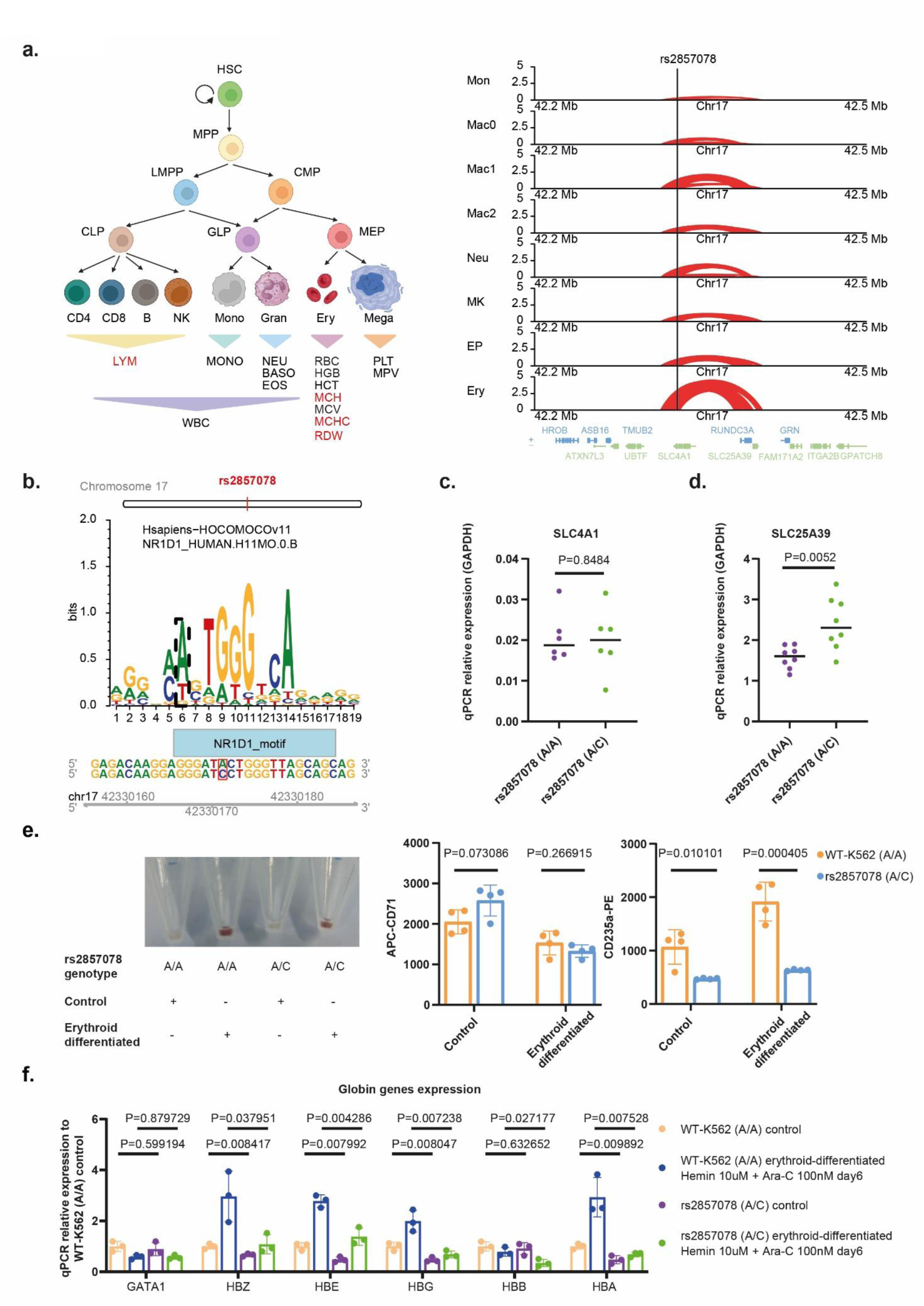
Functional profiling of silencer variant rs2857078 in hematopoiesis. **(a)** Left panel: schematic of the human hematopoietic hierarchy with blood cell traits assigned to their corresponding lineages or cell types. Traits for which rs2857078 was fine-mapped as a putative causal variant are highlighted in red (LYM: PIP = 0.38; MCH: PIP = 0.40; RDW: PIP = 0.79; MCHC: PIP = 0.03). PIP, posterior inclusion probability (scale 0–1). Right panel: Normalized chromatin interaction strength between rs2857078 and the promoter regions of nearby genes across primary blood cell types. Mon, monocytes; Mac0–2, macrophages M0, M1, and M2; Neu, neutrophils; MK, megakaryocytes; EP, erythroid progenitor; Ery, erythroblasts. **(b)** Prediction of the disruptiveness of rs2857078 on the transcription factor NR1D1 binding motif using motifbreakR. **(c)** Quantification of the effect of rs2857078 on SLC4A1 expression. SLC4A1 transcription was measured by qPCR in WT-K562 and rs2857078-KI cells (n = 6 biologically independent samples; expression relative to the housekeeping gene GAPDH is shown as mean ± s.d.; P values were calculated using a two-tailed unpaired t-test). **(d)** Quantification of the effect of rs2857078 on SLC25A39 expression. SLC25A39 transcription was measured by qPCR in WT-K562 and rs2857078-KI cells (n = 8 biologically independent samples; expression relative to the housekeeping gene GAPDH is shown as mean ± s.d.; P values were calculated using a two-tailed unpaired t-test). **(e)** Quantification of the effect of rs2857078 on erythroid differentiation efficiency by FACS. To induce erythroid differentiation, WT-K562 and rs2857078-KI cells were treated with 10 μM hemin and 100 nM Ara-C for 6 days. Cell pellets are shown before and after differentiation. For FACS analysis, cells were stained with CD71-APC and CD235a-PE antibodies, and the geometric mean fluorescence intensity of both markers was quantified (n = 4 biologically independent samples; values are shown as mean ± s.d.; P values were calculated using a two-tailed unpaired t-test). **(f)** Quantification of the effect of rs2857078 on globin gene transcription. Globin gene transcription was measured by qPCR in WT-K562 and rs2857078-KI cells before and after erythroid differentiation (n = 3 biologically independent samples; expression relative to WT-K562 control cells is shown as mean ± s.d.; P values were calculated using a two-tailed unpaired t-test).

To assess the endogenous functional consequence of *rs2857078*, we introduced the alternative C allele into K562 cells using CRISPR-mediated knock-in. Following single-cell cloning and genotyping, one heterozygous A/C clone was selected for downstream analyses (**Supplementary Fig. 4b**). Compared with reference-genotype cells, the edited clone showed no significant change in *SLC4A1* expression by qPCR (**Fig. 4c**), whereas expression of the distal chromatin-interacting gene *SLC25A39* was significantly increased (**Fig. 4d**). RNA-seq provided concordant evidence of increased *SLC25A39* expression in the heterozygous A/C clone (**Supplementary Fig. 4c**). The established roles of SLC25A39 in mitochondrial glutathione import, iron homeostasis and erythroid development make it a biologically plausible effector gene at this locus ^59,60^. Additional nearby genes implicated by cis-eQTL analysis, including *ASB16-AS1, TMEM101, TMUB2 and C17orf53*, also showed increased expression in the heterozygous A/C clone by qPCR (**Supplementary Fig. 4d**). The coordinate expression change of multiple genes within this locus suggests that this silencer element harboring *rs2857078* may function as a core regulatory hub governing transcriptional activity at this locus. Among these genes, C17orf53 encodes a DNA interstrand crosslink repair factor^61^, providing another potential connection to cellular processes relevant to hematopoietic maintenance^62^. Whether these genes contribute individually or jointly to erythroid phenotypes requires further investigation.

We next examined *rs4808806*, a LoSF variant included in fine-mapped signals for RDW (PP = 0.02) and hematocrit (HCT; PP = 0.018)^28^ (**Supplementary Fig. 5a**). Analysis of ATAC-seq profiles from 13 primary blood cell types^31,63^ placed *rs4808806* within an intron of *ELL* in a region preferentially accessible in granulocyte–monocyte progenitors and monocytes (**Supplementary Fig. 5b**). Following CRISPR-mediated introduction of the alternative G allele into K562 cells, RNA-seq showed increased ELL expression (adjusted P-value= 2.59×10^-5^) (**Supplementary Fig. 5c**), Whole-blood cis-eQTL data independently associated *rs4808806* with ELL expression (P = 3.6 × 10^-23^). ELL encodes an RNA polymerase II transcription elongation factor^64^. and is a component of the super elongation complex, linking this candidate target gene to transcriptional programs involved in cell-state regulation, including hematopoiesis^65^. Integration of GWAS and eQTL data using summary-data-based Mendelian randomization (SMR) further prioritized *ELL* as a candidate gene associated with hematocrit (P_SMR_=1.96×10^-7^) (**Supplementary Fig. 5e**)^66^. It is possible that upregulation of *ELL* gene may influence the maturation of red blood cells^67,68^, as in the multi-ancestry GWAS meta-analysis, the G allele of rs4808806 was associated with increased hematocrit (β = 0.0098, P = 1.90 × 10^-8^). Together, these results suggest that *ELL* as a potential mediator of the association at this locus. RNA-seq also detected upregulation in genes *CERS1* and *HOMER3* surrounding this *rs4808806* locus mediated by alternative G allele (**Supplementary Fig. 5c**) while blood-lineage promoter-capture Hi-C data supported contacts between the variant-containing region and *CERS1* and *HOMER3*^30^ (**Supplementary Fig. 5d**). In addition, rs4808806 was predicted to disrupt a motif for ZNF436, a reported transcriptional repressor (**Supplementary Fig. 5f**)^69^, placing ZNF436 as a candidate regulator of the allelic silencer effect.

Together, these two examples connect silencer-altering variants identified by screening to endogenous gene expression changes and indicate candidate target genes through complementary chromatin-interaction and genetic evidence. The expression changes involving multiple nearby genes suggest that the consequences of altered silencer activity may extend beyond a single target. These findings provide a basis for testing how silencer-mediated regulation contributes to blood cell trait variation.

### The silencer loss-of-function variant rs2857078 alters erythroid phenotypes in K562 cells

To investigate the phenotypic consequences of the silencer-disruptive variant rs2857078, we used K562 cells, an established model of induced erythroid differentiation in vitro^70–73^. Wild-type K562 cells (K562-WT) and a monoclonal knock-in line carrying the heterozygous rs2857078 A/C genotype (rs2857078-KI) were treated with 10 μM hemin and 100 nM cytarabine (Ara-C) for 6 days^73–75^. Following treatment, centrifuged cell pellets appeared red, consistent with hemoglobinization (**Fig. 4e**). We assessed the erythroid phenotype by flow cytometry for CD235a (glycophorin A), an erythroid surface marker, and CD71 (transferrin receptor 1), which mediates transferrin-dependent iron uptake^76^. After 6 days of differentiation, rs2857078-KI cells showed significantly lower CD235a surface expression than K562-WT cells, whereas CD71 expression did not differ significantly (**Fig. 4f and Supplementary Fig. 6**). These findings indicate an altered erythroid phenotype in the edited clone. The direction of this difference is compatible with the association of the rs2857078 C allele with lower hematocrit in the multi-ancestry GWAS meta-analysis (β = −0.0289, P = 9.67 × 10^-51^)^28^.

To further characterize the functional consequences of *rs2857078* on erythroid gene expression, we performed qPCR analysis of hemoglobin gene transcription following hemin- and Ara-C-induced erythroid differentiation. In K562-WT cells, the expression of embryonic and fetal hemoglobin genes *HBZ*, *HBE*, *HBG* and *HBA* increased significantly upon differentiation, whereas *HBB* expression remained unchanged (**Fig. 4g**). Although rs2857078-KI cells showed a similar overall pattern of globin induction, their post-differentiation expression levels of *HBZ*, *HBE*, *HBG* and *HBA* were significantly lower than those in K562-WT cells (**Fig. 4g**). These results demonstrate reduced globin transcript abundance in the edited clone and suggest a potential effect on hemoglobin production. The expression differences are consistent with the associations of the C allele with lower mean corpuscular hemoglobin (MCH; β = −0.0282, P = 7.88 × 10^-58^) and mean corpuscular hemoglobin concentration (MCHC; β = −0.0565, P = 5.46 × 10^-264^) in the multi-ancestry meta-analysis^28^. Together, the reduced CD235a expression and globin transcript abundance in rs2857078-KI cells support a role for this silencer-associated locus in erythroid regulation. These cellular findings provide a potential functional connection to the blood trait associations at the locus.

### Prioritizing silencer-altering variants using hematopoietic sequence-to-function models

Statistical fine-mapping prioritizes candidate causal variants at GWAS loci, but its resolution is often limited by linkage disequilibrium. Many silencer-altering variants identified in our screen had low posterior inclusion probabilities (**Supplementary Fig. 7a**), highlighting the value of complementary approaches to assess their regulatory potential. Recent sequence-to-function models predict molecular features from DNA sequence, including chromatin accessibility^32^ and gene expression^77^, providing a new way to investigate context-dependent effects of non-coding variation. To characterize these effects across hematopoietic populations, we trained ChromBPNet models using single-cell ATAC-seq data from nine populations: HSC, MPP, LMPP, GMP, CMP, CLP, MEP, pDC and monocytes^3278^. The models can predict the base-resolution of read count distribution of cell-type-specific ATAC-seq profiles from local DNA sequence contexts around peak regions^32^. We trained five-fold cross-validated models for each cell type (n=45 models in total). Predicted accessibility counts within peak regions correlated with observed counts on held-out chromosomes, with a median Pearson correlation of 0.698 (**Supplementary Fig. 7b**). Interpretation using DeepLIFT and TF-MoDISco^7980^, recovered motifs associated with hematopoietic lineage regulators (**Supplementary Fig. 7c**), supporting the use of these models to investigate sequence features underlying cell-type-specific accessibility.

We next integrated predicted regulatory effects with evolutionary constraint using a lasso regression framework based on FLARE^33^. The model used cell-type-specific ChromBPNet variant-effect predictions, accessibility annotations and local gene constraint features to predict phyloP scores. This approach estimates the component of evolutionary constraint captured by the selected regulatory features and generates a score for variant prioritization. The relationship between measured conservation and predicted regulatory effects supported integrating these complementary sources of information (**Fig. 5a**). We ranked variants by predicted phyloP, using these scores to prioritize candidates for functional interpretation rather than to assign probabilities of trait causality or pathogenicity. The analysis included 9,857,211 non-coding variants from the 1000 Genomes Project Phase 3 reference panel: 5,201,423 previously curated common variants with minor allele frequency greater than 5%^33^ and 4,655,788 low-frequency or rare variants in European populations^51,52^. The model incorporating quantitative variant-effect predictions across all nine populations (9CT-Full) outperformed the accessibility-overlap baseline (9CT-PeakOverlap) in predicting phyloP on held-out chromosomes, both genome-wide (P = 0.0012) and within accessible regions (P = 9.24 × 10^-5^) (**Supplementary Fig. 7d**). Prominent selected features included interactions between peak overlap and ChromBPNet-predicted log fold changes (**Fig. 5b**), indicating that quantitative sequence-based predictions provided information beyond accessibility overlap alone. Coefficients differed across hematopoietic populations, with prominent contributions from early progenitor contexts. These differences describe the predictive contributions of the features within the fitted model.

**Figure 5.**
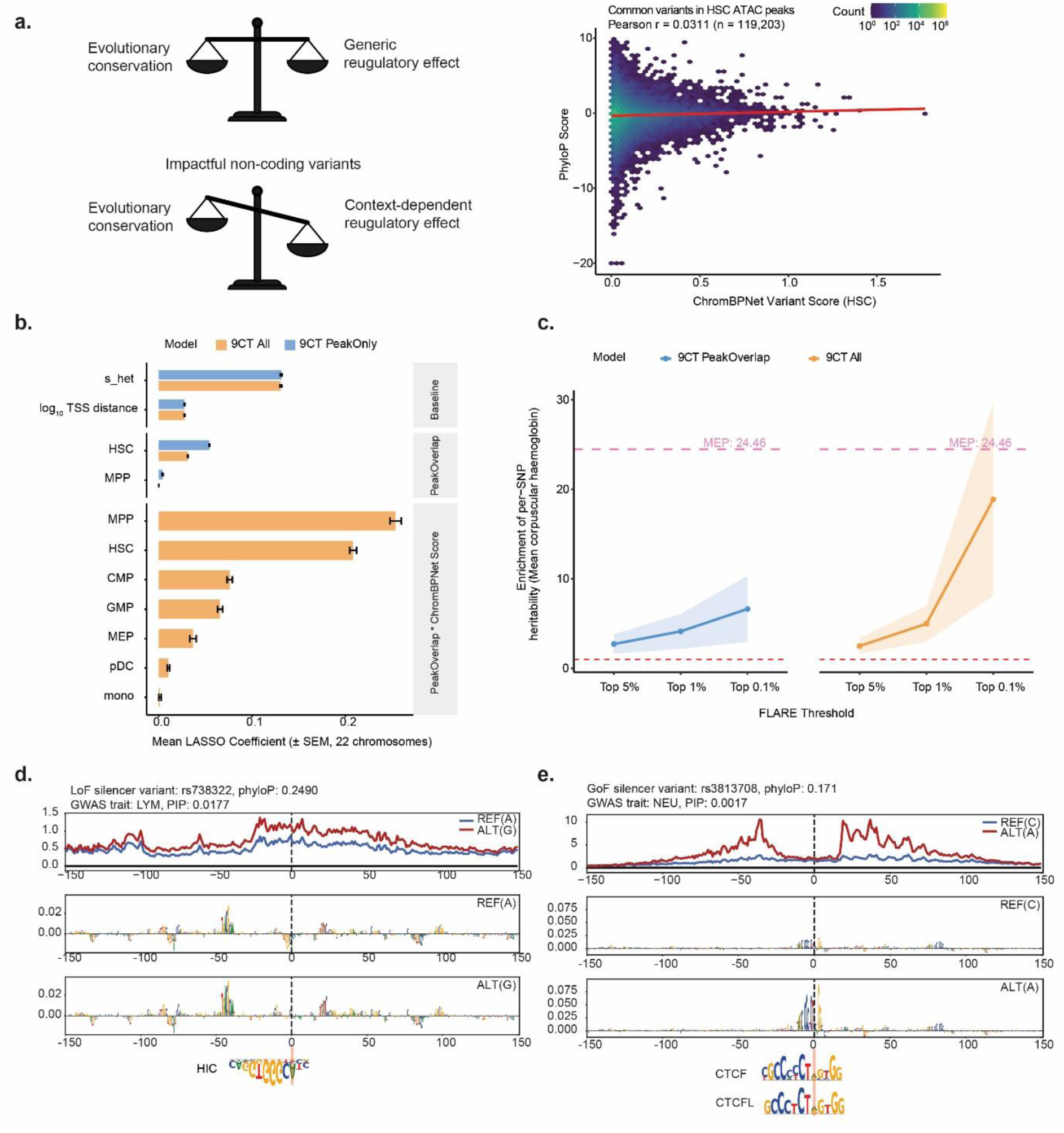
A machine learning framework to prioritize functional silencer variants. **(a)** Schematic illustrating the relationship between evolutionary conservation and the regulatory effect of non-coding variants. Deleterious non-coding variants that exert strong, context-specific regulatory effects are expected to be under stronger evolutionary constraint. Right: scatterplot of evolutionary constraint (PhyloP score) versus HSC-specific regulatory potential (ChromBPNet prediction) for common variants from the 1000 Genomes Project reference panel. **(b)** Feature importance in the LASSO regression. Horizontal bars show mean LASSO coefficients across 22 leave-one-chromosome-out folds for each feature selected in at least one fold. Features are grouped into four categories: Baseline (gene distance, s_het), Peak Overlap (binary peak overlap per cell type), and Interaction (peak overlap × log2FC). Within each group, features are ordered by mean coefficient in the 9CT-All model. Orange bars represent the 9CT-All model (29 features); blue bars represent the 9CT-PeakOnly model (11 features, peak overlap only). Error bars indicate ± s.e.m. across the 22 chromosomes. Features never selected (coefficient = 0 in all 22 folds) in either model are omitted. Bar magnitude reflects each feature’s contribution to predicting PhyloP conservation scores. **(c)** S-LDSC enrichment of per-SNP heritability for the blood cell trait MCH in prioritized variants. Variants were ranked by predicted PhyloP conservation score and stratified into three thresholds: top 5%, top 1%, and top 0.1%. Two FLARE models are compared across facet panels (colors as indicated). Lines connect mean enrichment estimates across thresholds; shaded ribbons indicate 95% confidence intervals (±1.96 × s.e.m.). The red dashed horizontal line at 1 indicates the baseline (no enrichment). Pink dashed horizontal lines indicate the enrichment of the biologically expected progenitor cell type, MEP, for trait MCH. **(d, e)** ChromBPNet predictions for two prioritized impactful silencer variants. Predictions are shown for a 300-bp window surrounding each variant, with predicted profiles for the reference and alternate alleles shown in blue and red, respectively. (**d**) The identified LoSF variant rs738322 is predicted to disrupt the motif of the transcriptional repressors HIC1/HIC2, increasing chromatin accessibility in K562 cells. (**e**) The identified GoSF variant rs3813708 is predicted to disrupt the CTCF/CTCFL motif, decreasing chromatin accessibility in K562 cells.

To assess the relationship between model-based prioritization and blood cell traits, we used stratified LD score regression to evaluate annotations comprising the top 5%, 1% and 0.1% of variants ranked by predicted phyloP (**Fig. 5c, Supplementary Fig. 7e-g**). After accounting for the baseline model, FLARE-prioritized variant annotations showed enrichment for blood cell trait heritability, with estimates varying across prioritization thresholds. For MCH, variants in the top 0.1% had a per-SNP heritability enrichment estimate comparable to that of variants within MEP-specific ATAC-seq peaks, although the estimate had substantial uncertainty (**Fig. 5c**). We also compared CADD scores across prioritized variants within and outside accessible regions^81^ (**Supplementary Fig. 7h-i**). Thes results suggest that FLARE prioritization captures regulatory variation relevant to blood cell traits. Prioritized variants within accessible regions also showed higher CADD scores than those outside accessible regions, consistent with greater predicted functional impact (**Supplementary Fig. 7h-i**).

Intersecting the top 0.1% of model-ranked variants with the experimentally identified silencer-altering variants yielded 12 candidates, of which 10 had PIPs below 0.1. We examined two examples by combining predicted allelic accessibility profiles, motif changes and eQTL associations (**Fig. 5d-e**). The LoSF variant rs738322 had a PIP of 0.0177 for lymphocyte count and was predicted to disrupt a HIC1/HIC2 repressor motif. Its eQTL associations implicated increased expression of PLA2G6 and MAFF, suggesting candidate links between the observed loss of repression and local gene regulation. PLA2G6 encodes calcium-independent phospholipase A₂β, which participates in phospholipid metabolism^82^. Previous work suggested that it plays a role in lymphocyte proliferation^83^. Thus, it is possible that upregulation of PLA2G6 by silencer variant rs738322 may increase lymphocyte count by enhancing proliferative capacity. On the other hand, the GoSF variant rs3813708 had a PIP of 0.0017 for neutrophil count and was predicted to alter CTCF/CTCFL motifs. Its eQTL associations implicated reduced expression of CCT6P3, GTF2IP14, ERV3-1 and ASL. Among these genes, ASL encodes argininosuccinate lyase, which contributes to arginine synthesis and nitric oxide production^84^. The involvement of arginine and nitric oxide metabolism in myeloid biology⁸⁵ suggests a possible connection to the associated trait^85^.

Together, these analyses integrate experimentally measured silencer activity with hematopoietic sequence-based predictions to nominate variants and regulatory mechanisms for further investigation. The identification of high-scoring silencer-altering variants with low PIPs illustrates how functional and computational evidence can complement statistical fine-mapping without independently establishing trait causality.

## Discussion

Functional characterization of non-coding GWAS variants is essential for understanding the molecular basis of human trait variation and disease susceptibility. Much of this work has focused on promoters and enhancers, leaving the contribution of silencer-associated variation less well understood. Here, we systematically evaluated 14,720 non-coding variants associated with 15 blood cell traits using the ReSE screening system. In K562 cells, 805 variants (5.47% of those selected for screening) showed allele-dependent silencer activity, comprising 425 loss-of-silencer-function and 380 gain-of-silencer-function variants. Motif analyses nominated disruption or strengthening of transcription factor binding sites as potential mechanisms underlying these effects. Together, these findings identify silencer-associated variation as a component of blood trait-related regulatory diversity. The chromatin annotations and heritability enrichment patterns further connect these variants to hematopoietic regulatory contexts, although activity measured in K562 cells cannot represent the full diversity of primary blood cell states. Extending these analyses to additional hematopoietic populations and organs involved in systemic blood cell regulation could reveal further context-dependent effects.

ReSE measures repression using a lentivirally integrated reporter, providing a chromosomal assay context that differs from episomal approaches such as Ss-STARR-seq and Silencer-seq^15129^. Integration nevertheless places the tested sequences outside their native genomic locations, where their activity may differ because of local chromatin organization, promoter compatibility and transcription factor availability. We therefore combined screening results with eQTL and promoter-capture Hi-C data to nominate endogenous target genes. Concordance between reduced silencer activity and increased gene expression, or between increased silencer activity and reduced expression, provides supporting evidence for these assignments. However, eQTL associations do not independently establish the causal variant or regulatory mechanism. Endogenous editing at rs2857078 and rs4808806 provided additional evidence linking these loci to gene expression changes. At rs2857078, the edited K562 clone also showed reduced CD235a expression and globin transcript abundance following induced erythroid differentiation, suggesting a functional connection to erythroid biology. Establishing which target genes mediate these cellular effects, and whether the same mechanisms explain the associated blood traits, will require further perturbation and rescue experiments. The expression changes observed after editing, together with promoter-capture Hi-C contacts, also suggest that silencer-associated variants can influence genes beyond the nearest annotated gene. Changes involving multiple nearby transcripts raise the possibility that individual silencer elements affect more than one target, although direct regulation remains to be distinguished from secondary transcriptional responses. Combining functional screens with chromosome-conformation assays and targeted endogenous perturbations could help resolve these relationships. Promoter-capture Hi-C can nominate promoter contacts, whereas H3K27me3 HiChIP could investigate interactions involving the subset of candidate silencers associated with this chromatin mark. Such analyses would help connect silencer activity to the organization of endogenous regulatory domains.

Many screen-identified silencer-altering variants had low fine-mapping PIPs (**Supplementary Fig. 7a**), while annotations defined by these variants showed enrichment for blood cell trait heritability (**Fig. 3c**). These observations concern different levels of evidence: fine-mapping estimates support for individual variants, whereas partitioned heritability evaluates the collective contribution of an annotation. To provide complementary functional prioritization, we applied a FLARE-based framework that predicts evolutionary constraint from hematopoietic regulatory features^33^. High-scoring annotations showed blood-trait heritability enrichment, and comparisons with CADD provided additional evidence of predicted functional importance (**Fig. 6c** and **Supplementary Fig. 7e-i**). ChromBPNet interpretation further nominated allele-dependent motif changes in specific hematopoietic contexts. These predictions complement the experimental screen by identifying candidate regulatory mechanisms. Training and evaluating models directly against experimentally measured allelic silencer effects could extend this framework toward predicting silencer-altering activity at untested variants.

Several considerations guide interpretation of these findings and future applications of ReSE. First, reliable hit identification requires adequate library representation, biological replication and statistical control of multiple testing. Allelic comparisons must also account for differences in construct abundance and dependence among overlapping sequence windows. Second, regulatory activity depends on cellular context. Although K562 and HepG2 provide tractable screening systems, validation in primary cells and relevant differentiation states is needed to establish physiological relevance. Third, ReSE detects sequences that reduce inducible caspase-9 reporter activity sufficiently to confer a survival advantage under the assay conditions. It therefore measures repression in a defined reporter and selection context, rather than directly identifying the molecular mechanism of repression. Endogenous nucleotide editing can test the effects of individual variants, whereas targeted deletion can assess the contribution of the surrounding element.

Our study combines experimental screening, endogenous editing and computational prioritization to investigate silencer-associated variation at blood cell trait loci. By identifying allelic effects on repression and linking selected variants to endogenous gene expression and erythroid phenotypes, this work extends functional interpretation of GWAS loci beyond promoter- and enhancer-centered approaches.

## Methods

### Cell culture

K562 cells were cultured in RPMI 1640 + L-Glutamine (Gibco), 10% fetal bovine serum (Biowest), and 1% Penicillin-Streptomycin (Gibco). HepG2 cells were cultured in DMEM, 10% FBS and Antibiotic-Antimycotic. Cell density and culture conditions were maintained according to the ENCODE Cell Culture Guidelines.

### Library Design and Construction

The initial SNPs list were retrieved from a trans-ethnic meta-analysis of 15 blood cell traits (RBC count, hemoglobin, hematocrit, mean corpuscular volume, mean corpuscular hemoglobin, MCH concentration, RBC distribution width, WBC count, neutrophils, monocytes, lymphocytes, basophils, eosinophils, platelet count, and mean platelet) in 746,667 individuals from five different ancestries (European ancestries, South Asian ancestries, Hispanic ancestries, East Asian ancestries, and African ancestries) generated by the Blood Cell Consortium (BCX) consortium (https://www.mhi-humangenetics.org/en/resources/). All the SNPs from fine-mapped 95% credible sets for 15 blood traits in trans- and sub-ancestry were first merged and deduplicated. The unique GWAS variants list was annotated by VEP and only retained the SNPs falling into the noncoding regions by excluding the coding or splice site variants. For each variant, six 130-nt constructs were designed based on the hg19 reference genome, such that the SNP site was located at the 1/4 (left), 1/2 (middle), or 3/4 (right) position of the construct, with each position represented by both alleles of the SNP. For the ReSE_SNP oligo library, each insert carrying both SNP alleles was designed for compatibility with the screening lentiviral vector pLenti-FKBP-delCasp9-Puro^9^ (with some modifications at the cloning sites) in the format: ACACGACGCTCTTCCGATCT-[130-nt insert]-AGATCGGAAGAGCACACGTC. The oligo pools were then ordered from GenScript.

The ReSE screen lentivirus vector pLenti-FKBP-delCasp9-Puro^9^ was digested with BsmBI enzyme and gel-purified. The PCR amplified oligo library was then inserted into the digested plasmids, 15 bp upstream of the EF-1α promoter, using Gibson Assembly. Each 20 μl assembly reaction contained 50 ng of insert DNA, 50 ng of digested plasmid, and 10 μl of 2× Gibson Assembly Master Mix, and was incubated at 50 °C for 60 min. Subsequently, 2 μl of the assembly mix was electroporated into 25 μl of Endura electrocompetent cells to assess transformation efficiency. Electroporation was scaled up to yield approximately 160,000 colonies, which were plated across four 245-mm Petri dishes containing 100 μg ml^-1^ carbenicillin. Colonies were scraped, and plasmid DNA was extracted using the QIAGEN Maxiprep Kit.

### Lentivirus production and infection

The lentivirus production and infection were performed as previously described^9^. The 293T cells were grown in five T175 flasks to 50% confluency before transfection. For each flask of 293T cells, maintained in 25 ml of fresh medium, 15 μg of library plasmid, 10 μg of psPAX2, 5 μg of pCMV-VSV-G, and 90 μl of X-tremeGENE 9 DNA Transfection Reagent were mixed in 1 ml of serum-free medium and added to the cells. Fresh medium was replaced the day after transfection. Virus-containing supernatant was collected on days 2 and 3 post-transfection, pooled, and concentrated using Lenti-X according to the manufacturer’s protocol. Virus titer was then determined by serially diluting (10^-3^ to 10^-10^) 4 μl of frozen virus supernatant in medium containing 8 μg ml^-1^ of polybrene to infect 293T cells. Two days after infection, cells were selected with 2 μg ml^-1^ puromycin for an additional 7 days, and viral titer was calculated from the number of surviving colonies at each dilution. K562 cells and HepG2 cells were then infected with the same viral library at a multiplicity of infection (MOI) of 0.3 by spin-infection. For spin-infection, 3×10^6^ cells per well of a 12-well plate were infected in 1 ml of medium containing 8 μg ml^-1^ of polybrene; four plates were used per infection for a total of 1.5×10^8^ cells, which resulted in approximately 300× to 500× coverage of ReSE_SNP library. Two days after infection, cells were selected by 2 ug ml^-1^ of puromycin for a further 5 days. For each biological replicate in both K562 and HepG2 cells, the infection was repeated using lentivirus from the same pool, ensuring identical library content across replicates.

### ReSE silencer screen

The ReSE screen was performed as previously described^9^. After puromycin selection, an aliquot of 3.5×10^8^ K562 cells infected with the ReSE_SNP viral library was frozen as the control population (pre-dimerizer). A separate aliquot of 3.5×10^8^ cells was treated with 1 nM AP20187 (Takara Bio 635060) for 18 h to induce apoptosis (post-dimerizer), after which dead cells were removed using a Dead Cell Removal Kit (Miltenyi Biotec). In the K562 ReSE screen, approximately 45.6 % of cells were recovered as live relative to the original input cell number after 18 h of AP20187 treatment. Accounting for cell growth during this 18-h period, the true survival rate is around 27.1 % (given a normal doubling time of 24 h for K562 cells). Live cells were expanded for a further 5 days. Genomic DNA was isolated from 3.5×10^8^ cells of either the non-treated control or the post-apoptosis-induction samples using the QIAamp DNA Blood Maxi Kit, with two columns processed per treatment. For HepG2 cells, the procedure was similar, except that dead cells were removed by washing away the medium, as HepG2 cells are adherent and live cells remained attached to the tissue culture flasks.

The ReSE lentiviral inserts within genomic DNA were amplified by PCR using flanking primers bearing Illumina sequencing adaptors (**Supplementary Table**). Each 100-μl PCR reaction contained 10 μg of genomic DNA, 20 μl of 5× Phusion HF buffer, 2 μl of 10 mM deoxynucleotide triphosphates, 2.5 μl of Phusion polymerase, 5 μl of 25 μM Primer 1.0, and 5 μl of 25 μM Primer 2.0. For each treatment sample, 16 reactions were prepared and pooled. The PCR program was: 98 °C for 30 s; 20 cycles of 98 °C for 10 s, 65 °C for 30 s, and 72 °C for 30 s; and a final extension at 72 °C for 5 min. PCR products were size-selected and purified, and the final products were sequenced on an Illumina NovoSeq 6000 platform.

### Luciferase assay

Candidate silencer sequences containing SNPs were amplified with primers carrying a homologous arm from either the input plasmid library or genomic DNA of K562 cells. These fragments were then inserted upstream of the promoters of the luciferase plasmids pGL4.23 (Promega; modified at the cloning sites; used to detect enhancer activity) and pGL4.53 (Promega; used to detect silencer activity) using NEBuilder HiFi. When a fragment containing the alternative allele could not be obtained by PCR amplification, site-directed mutagenesis was performed; primers used to introduce the desired mutations into the template plasmid. Cells were co-transfected with the pRL-CMV Renilla reporter plasmid and either pGL4.53 or pGL4.23 carrying the inserted silencer sequences. Luciferase assays were performed using the Dual-Luciferase Reporter Assay System (Promega) according to the manufacturer’s protocol. The original luciferase plasmid without any insertion served as the control. All luciferase assays were performed with three independent transfections carried out on different days. Used primers are listed in the **Supplementary Table**.

### CRISPR-Cas9 knock-in risk alleles of SNP in K562

Single-guide RNAs targeting GWAS SNP loci were cloned into the CRISPR plasmid pSpCas9(BB)-2A-Puro (PX459) V2.0 (Addgene: 62988) and co-transfected into K562 cells along with single-stranded HDR donor DNA templates containing the risk allele, using either the Bio-Rad or the 4D-Nucleofector system. After 2 days of transfection, cells were selected with 2 ug ml^-1^ puromycin. Three days after selection, cells were seeded into fresh medium without puromycin. Expanded clones were seeded into 96-well plates, and genomic DNA was extracted for PCR amplification of the regions containing the SNPs. Heterozygous and homozygous knock-in clones were confirmed by Sanger sequencing. The gRNA sequences used are listed in the **Supplementary Table**.

### Erythroid differentiation of K562 cells

For erythroid differentiation, 3×10^5^ K562 cells were seeded per well of a 6-well plate in 3 ml of complete culture medium. Differentiation was induced by treatment with 10 μM hemin (Sigma-Aldrich, cat. no. H9039) and 100 nM cytarabine (Ara-C; Sigma-Aldrich, cat. no. C1768) for 6 days, with medium replaced as needed.

### Flow cytometry analysis

Approximately 10^6^ cells were harvested and washed twice with staining buffer (2% BSA in PBS). Cells were resuspended in 100 μl of staining buffer and incubated with 5 μl of PE-conjugated anti-human CD235a (glycophorin A) antibody (BioLegend, cat. no. 349105) and 5 μl of APC-conjugated anti-human CD71 (transferrin receptor 1) antibody (BioLegend, cat. no. 334107) for 30 min at 4 °C in the dark. Stained cells were washed twice with staining buffer, resuspended in PBS, and immediately acquired on a flow cytometer. Data were analyzed using FlowJo v10.10.1 (BD Biosciences).

### Quantitative PCR

Total RNA was extracted using the ISOLATE II RNA Mini Kit, including on-column DNase I digestion (Bioline, cat. no. BIO-52073). Complementary DNA was synthesized using SuperScript IV VILO Master Mix (Invitrogen, cat. no. 11756050). Real-time PCR was performed using the SensiFAST SYBR No-ROX Kit (Bioline, cat. no. BIO-98020) on a Bio-Rad CFX Opus 384 real-time PCR system. Expression of the housekeeping gene GAPDH served as the normalization control. All qPCR experiments were performed with multiple biological replicates, and P values were calculated using an unpaired two-tailed Student’s t-test. Primer sequences are listed in the **Supplementary Table**.

### ReSE screen analysis

Cutadapt v3.4^88^ was used to extract the unique 130-nt SNP-containing sequences (5′ adapter: ACACGACGCTCTTCCGATCT; 3′ adapter: AGATCGGAAGAGCACACGT). Trimmed reads were then mapped to indexed references generated by Bowtie2^89^ from the oligo library designs, with the SNP sites masked as ‘N’. A custom script was used to extract the SNP nucleotide identity from the CIGAR strings of the BAM files, assign reads to the respective alleles, and quantify their abundance. The resulting read-count tables were first median-normalized to adjust for differences in library size and read-count distribution. Read-count variance was then estimated, and a negative binomial model implemented in MAGeCK^40^ was used to test whether fragment abundance differed significantly between post-apoptosis-induction and control replicates. P values were calculated from the negative binomial model using a modified robust ranking aggregation algorithm, and FDR values were computed from the empirical permutation P values using the Benjamini-Hochberg procedure. Because fold enrichments are only semi-quantitative, fragments with FDR < 0.01 were considered significant hits for downstream analyses, and the list of silencers was sorted by FDR in ascending order.

### Allelic silencer activity analysis

To assess differential allelic silencer activity, we devised a novel statistical framework, termed "rank-difference RRA testing," to analyze the ReSE screen data (**Supplementary Fig.3**). Briefly, the ranking difference between the effect and non-effect SNP constructs was extracted from the RRA analysis performed by MAGeCK. These ranking differences were then sorted and converted to quantile ranks (percentiles). Under the null-distribution assumption of uniformity for the quantile ranks, the modified RRA method α-RRA was applied to assess the significance of the ranking difference between the two allele constructs. A permutation test was used to compute the P values assigned to the significance scores, and FDR values were derived from the empirical permutation P values using the Benjamini-Hochberg procedure for multiple-testing correction.

### Motif disruptiveness analysis

To evaluate the disruptive effects of SNPs on transcription factor (TF) binding sites, the R package motifbreakR v2^43^ was used to scan the SNPs against TF position weight matrices (PWMs) acquired from HOCOMOCO v11 database^90^. The average difference between alternative and reference alleles were plotted together with the normalized expression levels of the corresponding TF genes from RNA-seq.

### S-LDSC heritability enrichment analysis

Stratified LD score regression (S-LDSC)^51,52^ was used to quantify the heritability explained by specific subset of GWAS SNPs, using binary annotations that partition SNPs into regions of interest while accounting for linkage disequilibrium (LD). The baseline model (v2.2) was used to assess heritability enrichment relative to the basic genomic annotation. The following statistics were reported: (1) the proportion of SNP heritability explained (Prop(h^2^)), (2) the per-SNP-heritability enrichment (enrichment = proportion of h^2^/proportion of SNPs) and (3) the standard deviation of heritability enrichment.

### Promoter-capture Hi-C analysis

The score matrix of promoter-capture interaction frequencies across 17 primary blood cell types^31^ was called using CHiCAGO with a cutoff of score > 5. Interaction scores were normalized using z-score transformation across all cell types. Relative interaction strengths, represented by arch height, were visualized with the R package plotgardener^91^.

### RNA-seq analysis

RNA was isolated using ISOLATE II RNA Mini Kit (BIO-52073), and sequencing libraries were prepared with the Invitrogen Collibri Stranded RNA Library Preparation kit (A39003024), according to the manufacturer’s protocol. Whole transcriptome sequencing was performed on the Illumina NovaSeq 6000 platform.

Bulk RNA-seq samples were processed using the Snakemake pipeline (https://github.com/snakemake-workflows/rna-seq-star-deseq2). Briefly, adapters were trimmed from reads using Cutadapt v3.4^88^, and spliced transcripts were aligned to the reference genome (GRCh37, Ensembl release 75) with STAR, which was also used to quantify per-gene read counts^92^. Differential expression analysis was performed with DESeq2^93^, and P values were corrected using the Benjamini– Hochberg procedure for FDR control.

### Preprocessing of hematopoietic scATAC-seq

Pseudobulk alignment files of scATAC-seq data for 9 hematopoietic lineages^78^ were converted to tagAlign format with Tn5 offset correction. Cell-type-specific ATAC-seq peaks were called using MACS2^94^ with the parameters "--nomodel --shift −100 --extsize 200 -p 0.01", and ENCODE blacklist regions^95^ were subsequently filtered from the called peaks.

### Training ChromBPNet models for hematopoietic cells

Hematopoietic cell-type-specific ChromBPNet models^32^ were trained using pseudobulk ATAC-seq data from 9 populations (HSC, MPP, LMPP, CMP, CLP, GMP, MEP, pDC, and monocyte). Model training used a chromosome-held-out fivefold cross-validation scheme as previously described^33^, ensuring that each chromosome was included in the test set in at least one fold. GC-matched non-peak background regions were generated as negative instances. To correct for Tn5 transposase sequence bias, a pre-trained K562 bias model was supplied during training of the cell-type-specific bias-factorized ChromBPNet models. Training proceeded for up to 50 epochs with early stopping if validation performance on held-out chromosomes ceased to improve, and the model with the lowest validation loss was saved.

### Discovering lineage-specific motif using DeepLIFT and TF-MoDISco

To identify lineage-specific motif instances predictive of chromatin accessibility from the cell-type-specific ChromBPNet models, we first generated base-resolution contribution scores in the filtered peak regions for each cross-validation-fold ChromBPNet model. Contribution scores were computed with the DeepLIFT algorithm implemented by DeepSHAP^96^, using 20 dinucleotide-shuffled reference sequences for each peak sequence. The mean contribution scores across the five folds were then computed for each cell type and used as input to run TF-MoDISco^80^ with the parameters “-w 400 -n 1000000”. TF-MoDISco identified a unique set of highly important sequence patterns ("seqlets") for each cell type, which were predictive of chromatin accessibility and matched known or novel TF binding sites learned by the cell-type-specific ChromBPNet models. The MotifCompendium tool was used to generate a non-redundant set of clustered patterns across all cell types and to annotate known motifs within the clustered patterns by similarity comparison. In total, 141 non-redundant clustered motifs were identified and annotated using a similarity score threshold of 0.96. The Fi-NeMo tool (https://github.com/kundajelab/Fi-NeMo) was then run against the consensus motif instances in the clustered compendium for each cell type to identify hits within the cell-type-specific peak regions. The occurrence matrix of motif instances in the compendium across all cell types was used for clustering analysis.

### Predicting cell-type-specific variant effects using ChromBPNet

To score the regulatory effects of 9,857,211 noncoding variants from the 1000 Genomes Project Phase 3 reference panel, the ChromBPNet models of each cross-validation fold were run for each cell type to predict allele-specific base-resolution scATAC-seq pseudobulk coverage profiles for the 1-kb genomic sequence centered on each variant (https://github.com/kundajelab/variant-scorer). The effect size of each variant was calculated as the log2 fold change in total predicted coverage within each 1-kb window for the alternate versus the reference allele, and was further aggregated as the mean of the log2 fold changes across the five folds for each cell type; this quantity was referred to as the ChromBPNet score.

### FLARE analysis in hematopoietic context

The FLARE analysis was performed as previously described^33^. Briefly, a lasso regression model was trained to predict evolutionary conservation from cell-type-specific ChromBPNet scores and local genomic features (that is, distance to the nearest TSS and the s_het_ constraint score of the nearest gene). The phyloP score, which measures the evolutionary conservation of each variant, was extracted from the phyloP100way track deposited in the UCSC Genome Browser. The nearest-gene s_het_ constraint score was calculated with GeneBayes^97^ as the selection coefficient against heterozygous loss-of-function variants for each protein-coding gene. FLARE models were trained under two schemes: (1) 9CT-PeakOverlap, predicting the phyloP score using features including the log10-transformed distance to the nearest TSS, the s_het_ constraint score of the nearest gene, and binary annotations of variants overlapping accessible regions in each given cell type; and (2) 9CT-Full, predicting the phyloP score using the same features plus the predicted accessibility effects (that is, the ChromBPNet score) in each given cell type. The coefficients of all features were constrained to be non-negative for biological interpretability. Models were trained using a leave-one-chromosome-out approach, resulting in 22 independent models, one for each chromosome. The optimal regularization parameter was determined via fourfold cross-validation, and model performance was evaluated on held-out chromosomes by comparing predictions with observed phyloP values using Pearson correlation.

### Heritability and CADD enrichment analysis for prioritized variants

Variants in the top 0.1%, 1%, and 5% ranked by predicted phyloP values were extracted for heritability and CADD enrichment analysis. S-LDSC^52^ was used to evaluate the per-SNP heritability enrichment of each blood cell trait for the binary annotation of prioritized variants at each threshold. The baseline and trait-corresponding blood-cell-type region-based heritability enrichments are indicated by dashed lines. CADD scores for each variant were extracted from the CADD v1.6 track^81^ on the UCSC Genome Browser. Odds ratios (ORs) were computed using a one-sided Fisher’s exact test, comparing the proportion of prioritized variants exceeding CADD PHRED thresholds of >10 and >20 against the full background set at each threshold.

## Data availability

The raw sequencing data including ReSE screen and RNA-seq generated in the study were deposited at GEO under accession code GSE347432. (reviewer access token:)

## Code availability

The scripts for reproducing the data analysis are available on GitHub at (https://github.com/MorganTan95/ReSE_SNP_paper).

## ACKNOWLEDGEMENTS

This work was supported by the Gisela Thier Fellowship from Leiden University Medical Center and the ERC Starting Grant 950655-Silencer from the European Research Council (all awarded to B.P.).

## AUTHOR CONTRIBUTION

B.P. conceptualized the project. M.T., H.V.W., and B.P. conducted investigations. M.T. conducted formal analysis. M.T. and B.P. wrote the original draft. B.P. supervised the project.

## Notes

### Competing Interest Statement

The authors have declared no competing interest.

### Summary of Updates

Figure 1-5 revised; Supplemental files updated

