## Supplementary Figure for "Genetic variants within silencer elements shape human blood cell phenotype diversity"

**Supplementary Figure 1**

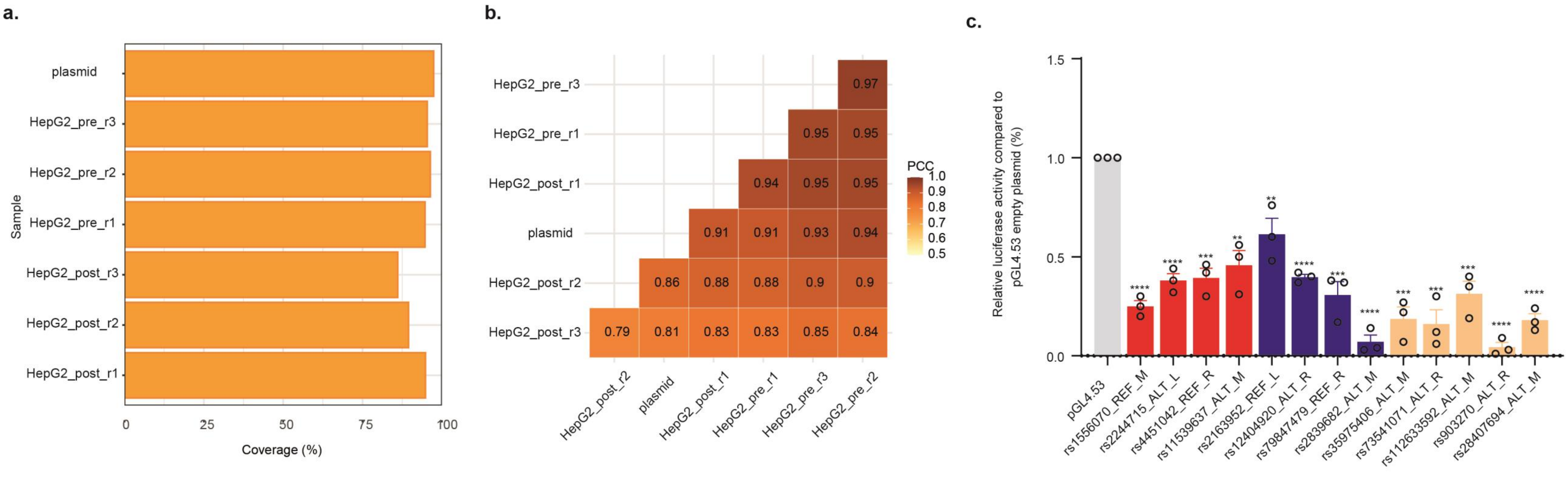


**Supplementary Figure 1. ReSE silencer screen at fine-mapped SNPs loci in HepG2 cells.**
(a) Coverage of ReSE SNP library complexity in the plasmid library input and in 3 biological replicates of screening samples (both pre- and post-dimerizer treatment) in K562 cells.
(b) Pearson correlation between biological replicates of the ReSE SNP libraries isolated from K562 cells (pre: cell population before dimerizer treatment; post: surviving cells after dimerizer addition; input: constructed ReSE plasmid library).
**(c)** Luciferase assays to validate the repressive activity of SNP-containing silencers identified from HepG2 cells with the cutoff of FDR < 0.01. Different colors indicate SNP-containing silencers from top (red) or bottom (blue) or random (orange) positions of the list ranked by FDR. Silencer regions were cloned by PCR from the input plasmid library DNA, then inserted upstream of the promoter of the luciferase reporter plasmid pGL4.53. and empty pGL4.53 plasmid was used as the control for baseline luciferase activity. The luciferase and renilla plasmid were co-transfected into HepG2 cells using electroporation. The y axis represents the percentage of luciferase activity compared to pGL4.53 empty plasmids in the respective cells (n = 3 biological independent samples; bars show mean value ± s.e.m.; $^{ns}P > 0.05$, $^{*}P < 0.05$, $^{**}P < 0.01$, $^{***}P < 0.001$, $^{****}P < 0.0001$, calculated using two-tailed unpaired t-test).

### Supplementart Figure 2

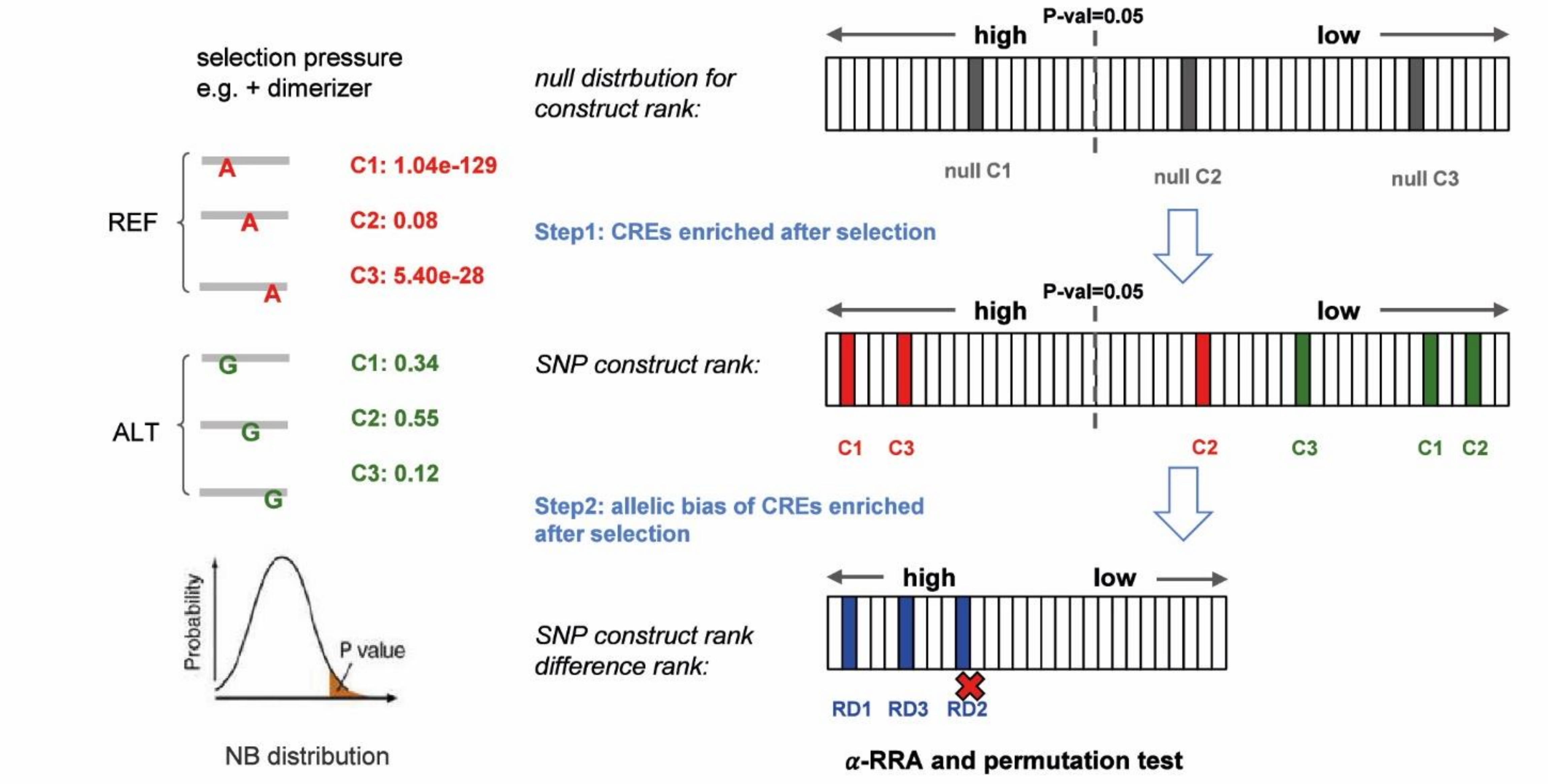


**Supplementary Figure 2. Overview of the rank-difference RRA framework for identifying silencer-associated genetic variants.**
Read counts from cis-regulatory element (CRE) constructs tiling each SNP, covering both the reference and alternate alleles, were first normalized. A mean-variance model was then fitted to the normalized counts from pre-dimerizer replicates to characterize baseline count variability. Differential activity between the two conditions (pre- vs. post-dimerizer) was assessed against a negative binomial null distribution, yielding one-sided P values reflecting positive selection. CRE constructs were ranked by these P values to produce a construct-level rank list. For each SNP tiled by six allelic CRE constructs (tiling positions: left, middle, right; alleles: reference and alternate), allelic bias was quantified as the difference in construct rank between the reference and alternate constructs at the same tiling position. These rank differences were independently ranked and normalized to percentiles, yielding a percentile score triplet (percentile_left, percentile_mid, percentile_right). Under the null hypothesis of no allelic difference, each percentile follows a uniform distribution, and the k-th order statistic of the three percentiles follows a Beta distribution. The allelic significance score ρ was defined as the minimum P value across the three order statistics, providing a robust summary of allelic imbalance. To reduce false positives, an α-robust rank aggregation (α-RRA) step was applied to exclude constructs with non-significant rank-difference scores ($P > 0.05$). Finally, an empirical false discovery rate (FDR) was assigned to each SNP's ρ score by permutation testing, in which the SNP construct rank list was shuffled to generate a null distribution of ρ values.

### Supplementary Figure 3

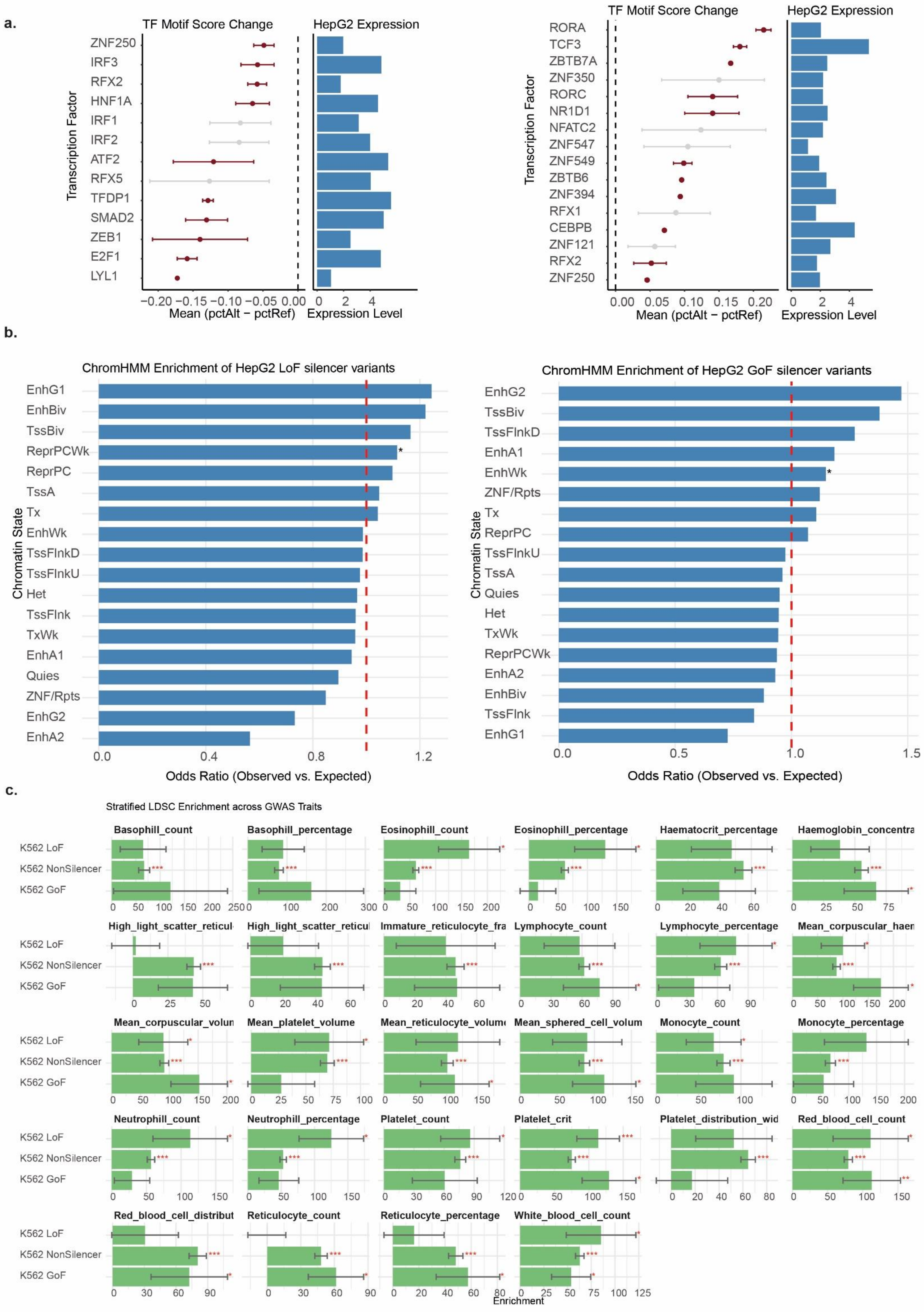

**Supplementary Figure 3. Epigenomic patterns and partitioned heritability of silencer-affecting GWAS variants**

(a) Disruptive TF binding sites associated with LoSF and GoSF variants in HepG2 cells. Left: schematic of how GWAS variants disrupt TF motifs, leading to LoF or GoF of silencer elements. Middle and right: recurrent disruptive TF motifs among LoSF (left) and GoSF (right) variants, scored and ranked with motifbreakR. The x axis shows the difference between the pctAlt and pctRef scores — the percentage of the best score each motif could achieve on an ideal reference or alternate sequence. TF motifs on the y axis are ranked by the mean pctAlt − pctRef difference; first and third quartiles of the difference are shown. RNA-seq expression levels of the TF genes in HepG2 cells, normalized as log(TPM + 1), are shown on the right.

(b) Chromatin states associated with functional transitions of silencer elements in HepG2 cells. Enrichment of ChromHMM states within LoSF or GoSF variants was calculated relative to a background of all SNPs included in the ReSE screen. The red dashed line indicates the odds-ratio cutoff, comparing the observed frequency of each of the 18 ChromHMM states among LoSF or GoSF variants with the expected frequency (background: all SNPs). P values were calculated using Fisher's exact test; *$P < 0.05$, **$P < 0.01$, ***$P < 0.001$.

(c) Enrichment of blood cell trait heritability in functional silencer variants identified in K562 cells. Heritability enrichment for blood cell traits associated with LoSF and GoSF variants identified in K562 cells was estimated with stratified LD score regression, which quantifies the heritability ($h^2$) enrichment of SNPs in each annotation (LoSF, GoSF, and non-silencer variants). Bar height represents the enrichment, calculated as the ratio of Prop_$h^2$ to Prop_SNP (Prop_$h^2$: proportion of total SNP heritability explained by the annotation; Prop_SNP: proportion of all SNPs in the study falling within the annotation). Error bars indicate the standard deviation of the enrichment. Asterisks indicate P values testing whether enrichment is significantly greater than 1. Trait abbreviations: basophil count/percentage, BASO/BASO%; eosinophil count/percentage, EOS/EOS%; hematocrit, HCT; hemoglobin concentration, HGB; high light-scatter reticulocyte count/percentage, HLR/HLR%; immature reticulocyte fraction, IRF; lymphocyte count/percentage, LYM/LYM%; mean corpuscular hemoglobin, MCH; mean corpuscular volume, MCV; mean platelet volume, MPV; mean reticulocyte volume, MRV; mean sphered cell volume, MSCV; monocyte count/percentage, MONO/MONO%; neutrophil count/percentage, NEUT/NEUT%; platelet count/plateletcrit, PLT/PCT; platelet distribution width, PDW; red blood cell count, RBC; red blood cell distribution width, RDW; reticulocyte count/percentage, RET/RET%; white blood cell count, WBC.

### Supplementary Figure 4

**a.**

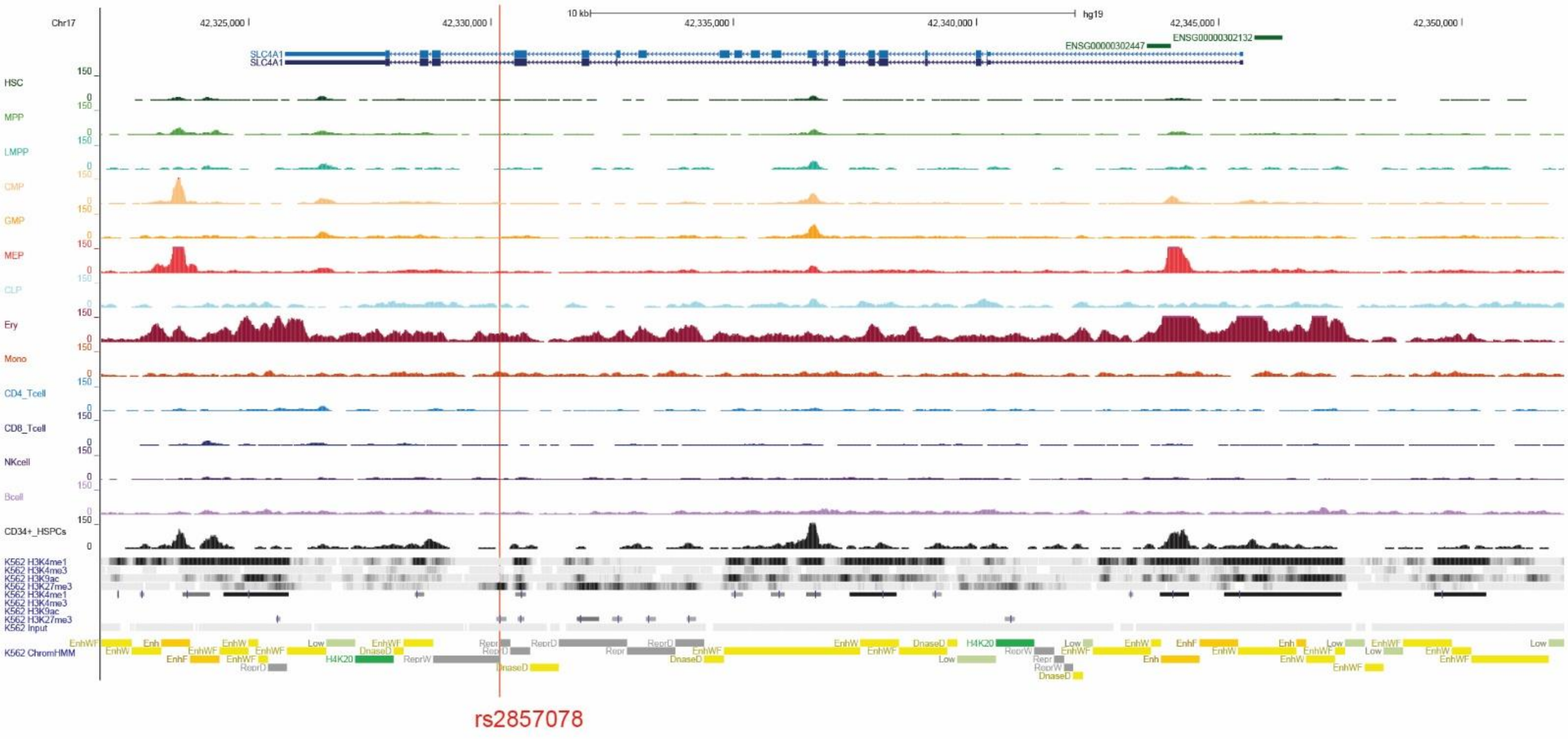


**b.**

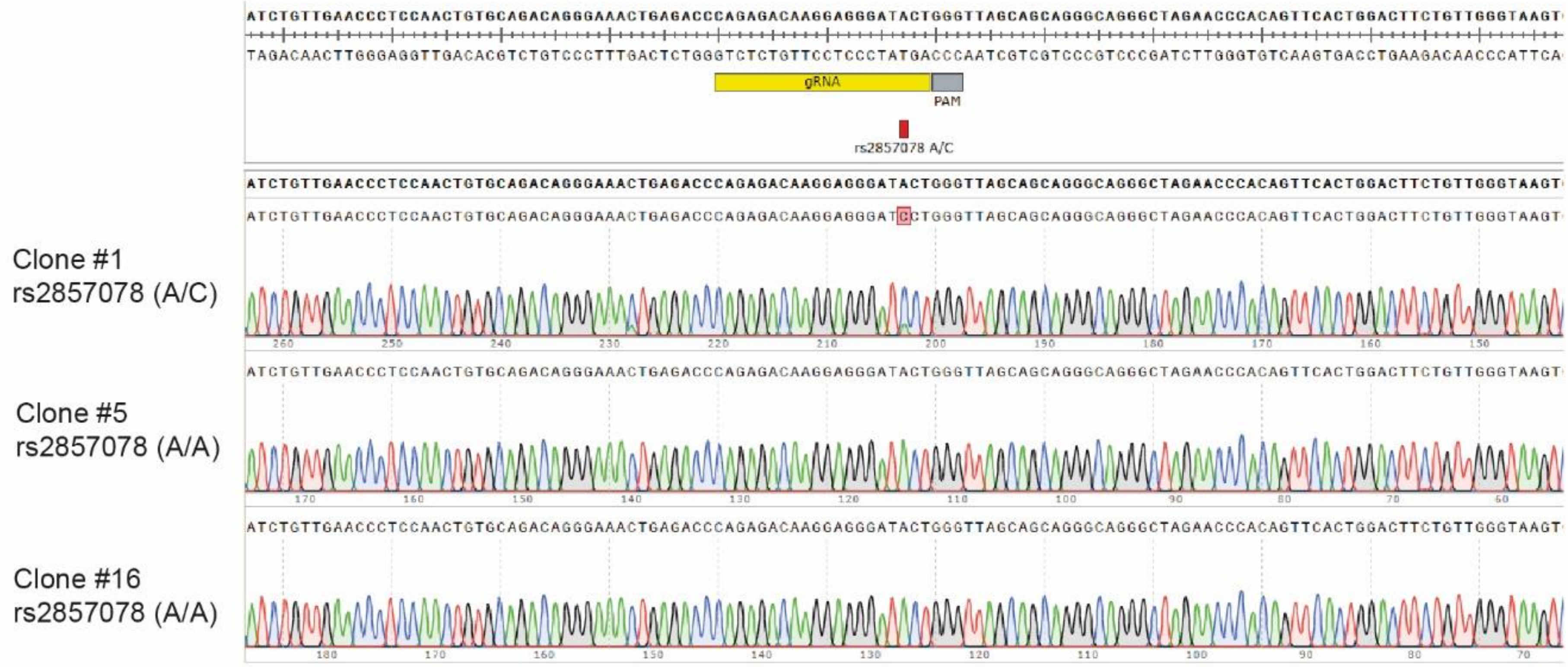


**c.**

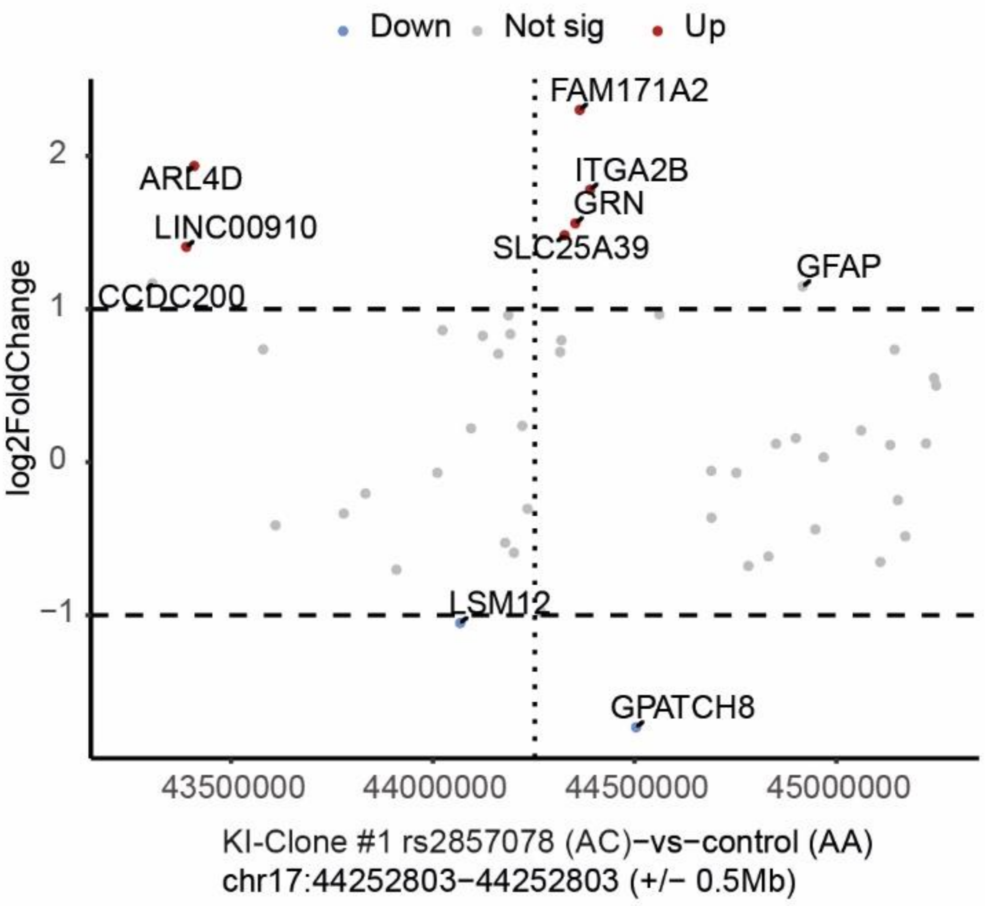


**d.**

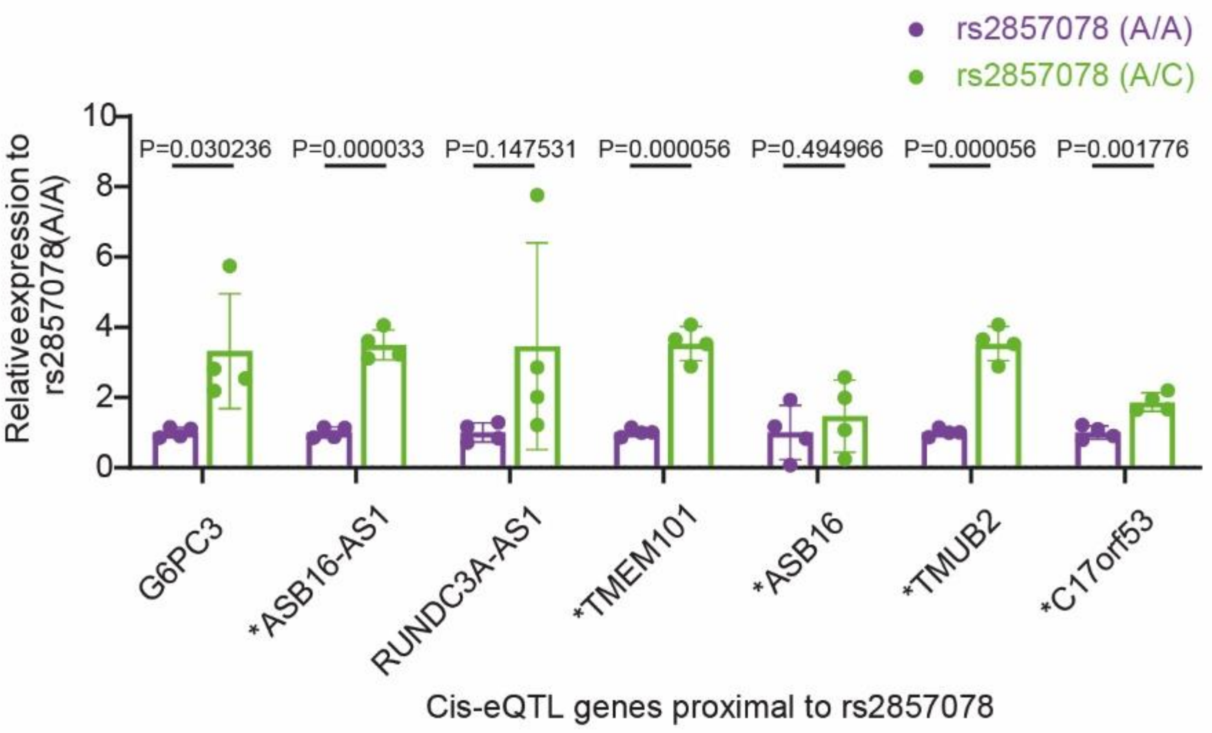

**Supplementary Figure 4. Linking the silencer-disruptive variant rs2857078 to its functional cellular context and target genes.**

(a) Normalized ATAC-seq profiles of 13 primary blood cell types at the rs2857078 locus.

(b) Sanger sequencing verification of the gRNA target region after CRISPR knock-in converting rs2857078 from A to C.

(c) RNA-seq expression changes of genes near rs2857078 after CRISPR knock-in. The x axis shows genomic coordinates of genes surrounding rs2857078 (−0.5 Mb to +0.5 Mb); the y axis shows log2 fold change determined with DESeq2. Dashed horizontal lines indicate the |log2FC| cutoff of 0.5, and the vertical dashed line indicates the position of rs2857078. Each dot represents one gene; red dots indicate significantly upregulated genes (FDR < 0.05 and log2FC > 0.5), blue dots indicate significantly downregulated genes (FDR < 0.05 and log2FC < −0.5), and gray dots indicate genes not meeting both thresholds.

(d) Quantification of the effect of rs2857078 on nearby gene expression. Cis-eQTL genes proximal to rs2857078 are highlighted. Gene transcription was measured by qPCR in WT-K562 and rs2857078-KI cells (n = 4 biologically independent samples; expression relative to the housekeeping gene GAPDH is shown as mean ± s.d.; P values were calculated using a two-tailed unpaired t-test).

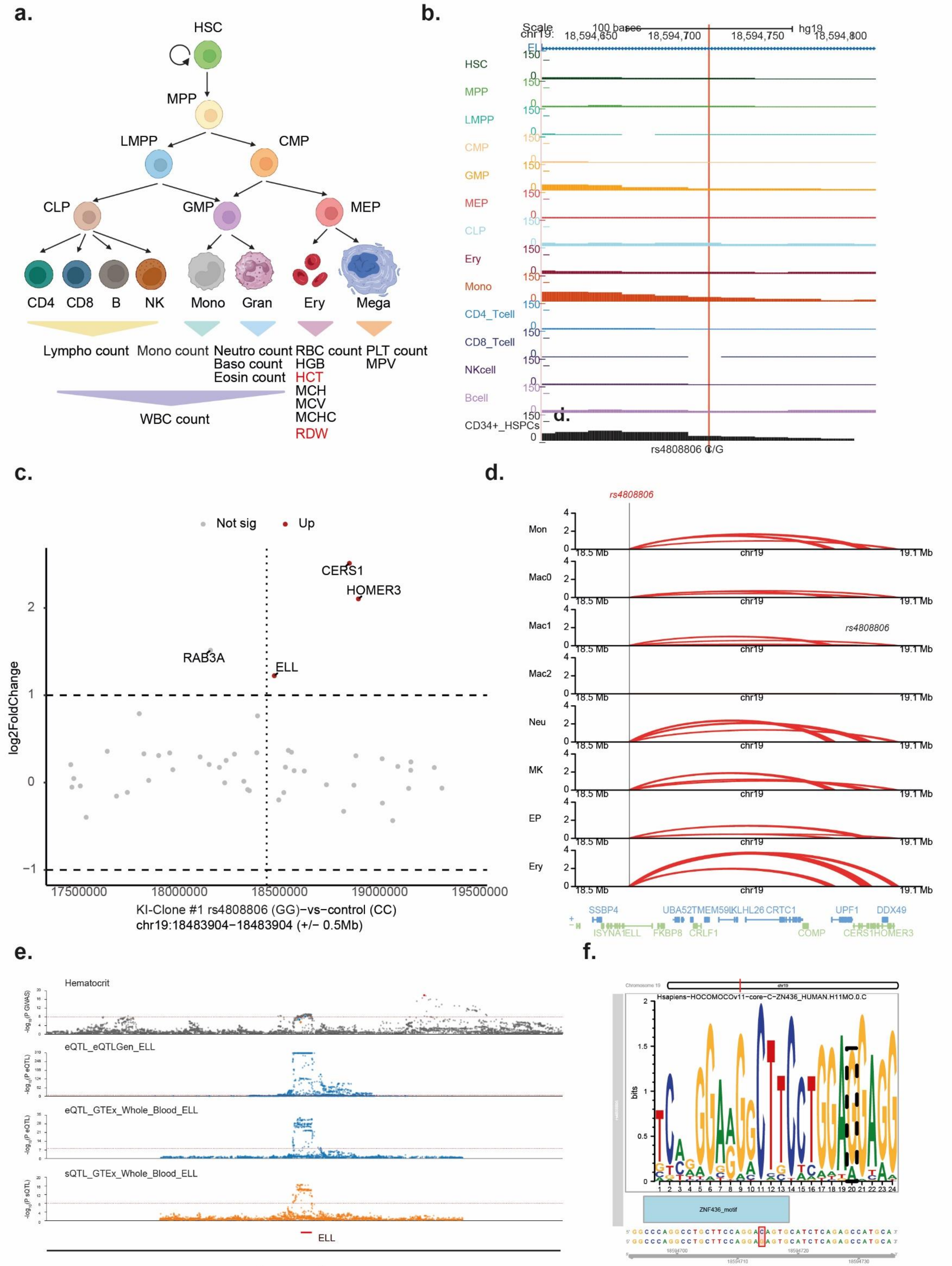
a.
HSC
MPP
LMPP
CMP
CLP
GMP
MEP
CD4
CD8
B
NK
Mono
Gran
Ery
Mega
Lympho count
Mono count
Neutro count
Baso count
Eosin count
RBC count
HGB
HCT
MCH
MCV
MCHC
RDW
PLT count
MPV
WBC count
b.
Scale
100 bases
hg19
chr19:
18,594,650
18,594,700
18,594,750
18,594,800
ELL
HSC
MPP
LMPP
CMP
GMP
MEP
CLP
Ery
Mono
CD4_Tcell
CD8_Tcell
NKcell
Bcell
CD34+_HSPCs
rs4808806 C/G
c.
Not sig
Up
CERS1
HOMER3
RAB3A
ELL
log2FoldChange
KI-Clone #1 rs4808806 (GG)−vs−control (CC)
chr19:18483904−18483904 (+/− 0.5Mb)
d.
rs4808806
Mon
Mac0
Mac1
Mac2
Neu
MK
EP
Ery
18.5 Mb
chr19
19.1 Mb
SSBP4
UBA52
TMEM59L
KLHL26
CRTC1
UPF1
DDX49
ISYNA1
ELL
FKBP8
CRLF1
COMP
CERS1
HOMER3
e.
Hematocrit
eQTL_eQTLGen_ELL
eQTL_GTEx_Whole_Blood_ELL
sQTL_GTEx_Whole_Blood_ELL
ELL
f.
Hsapiens-HOCOMOCOv11-core-C-ZN436_HUMAN.H11MO.0.C
bits
ZNF436_motif

**Supplementary Figure 5. Linking the silencer-disruptive variant rs4808806 to its functional cellular context and target genes.**

(a) Left: schematic of the human hematopoietic hierarchy (abbreviations as in Fig. 5a). Right: fine-mapping identifies rs4808806 as a putative causal variant for the blood cell traits HCT (hematocrit; PP = 0.018) and RDW (red blood cell distribution width; PP = 0.02).

(b) Normalized ATAC-seq profiles of 13 primary blood cell types at the rs4808806 locus.

(c) RNA-seq expression changes of genes near rs4808806 after CRISPR knock-in. The x axis shows genomic coordinates of genes surrounding rs4808806 (−0.5 Mb to +0.5 Mb); the y axis shows log2 fold change determined with DESeq2. Dashed horizontal lines indicate the |log2FC| cutoff of 0.5, and the vertical dashed line indicates the position of rs4808806. Each dot represents one gene; red dots indicate significantly upregulated genes (FDR < 0.05 and log2FC > 0.5), blue dots indicate significantly downregulated genes (FDR < 0.05 and log2FC < −0.5), and gray dots indicate genes not meeting both thresholds.

(d) Normalized chromatin interaction strength between rs4808806 and the promoter regions of nearby genes across primary blood cell types. Mon, monocytes; Neu, neutrophils; Mac0–2, macrophages M0, M1, and M2; EndP, endothelial precursors; MK, megakaryocytes; Ery, erythroblasts. Significant promoter-capture Hi-C loops linking rs4808806 to the genes UPF1, COPE, GDF1, UBA52, HOMER3-AS, and HOMER3 are shown; loop height indicates the relative interaction strength between rs4808806 and the target gene promoters in each lineage.

(e) Summary-data-based Mendelian randomization (SMR) analysis integrating the eQTL/sQTL signal of ELL with the GWAS signal for the hematocrit (HCT) trait.

(f) Prediction of the disruptiveness of rs4808806 on the transcription factor ZNF436 binding motif using motifbreakR.

### Supplementary Figure 6

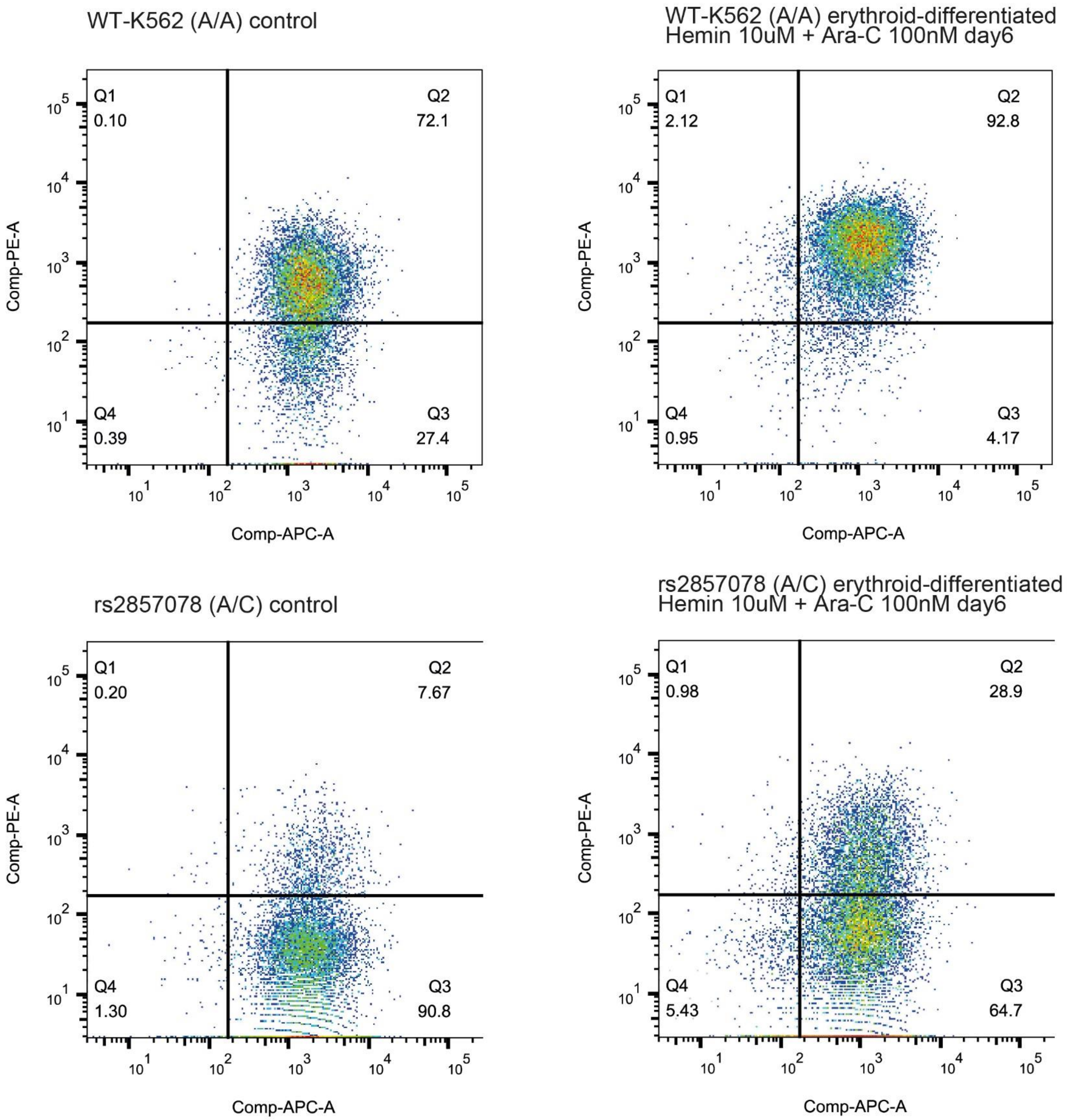


**Supplementary Figure 6. FACS gating examples quantifying the effect of rs2857078 on erythroid differentiation efficiency.** Representative flow cytometry plots of WT-K562 (A/A) and rs2857078-KI (A/C) cells under basal conditions (control) and after erythroid differentiation (10 μM hemin + 100 nM Ara-C, day 6). Cells were stained with CD71-APC (x axis) and CD235a-PE (y axis). Quadrant percentages are indicated; the upper-right quadrant (Q2, $CD71^{+}CD235a^{+}$) denotes erythroid-differentiated cells. Corresponding quantification across biological replicates is shown in Fig. 5f.

#### Supplementary Figure 7

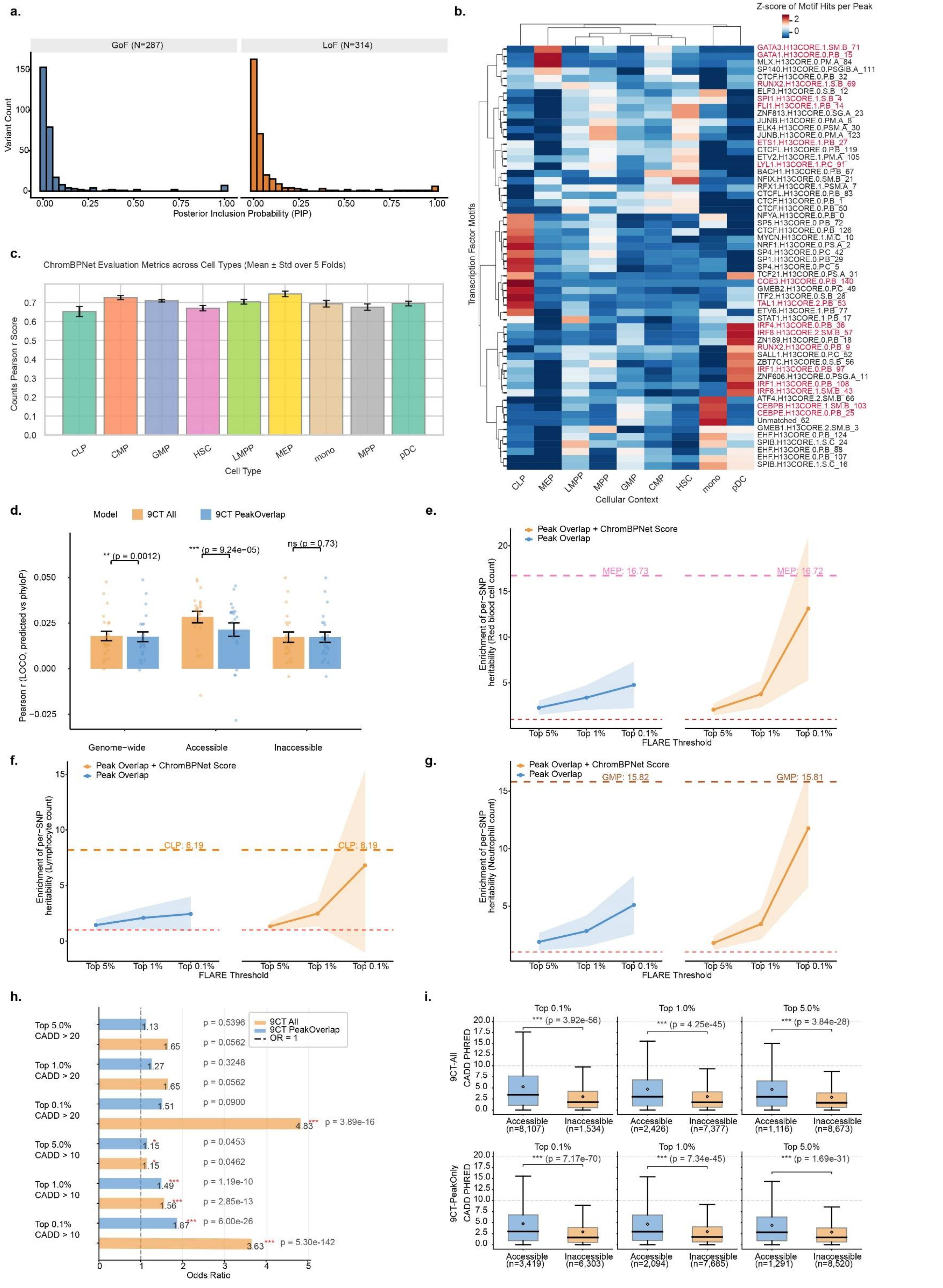

**Supplementary Figure 7. A machine learning framework to prioritize functional silencer variants.**
**(a)** Distribution of posterior inclusion probabilities (PIPs) for blood cell traits among GoSF and LoSF variants (defined in Fig. 6).
**(b)** Pearson correlation between predicted and observed chromatin accessibility profiles (total counts) in held-out chromosomes across all folds of the ChromBPNet models.
**(c)** Heatmap of motif instances identified by Fi-NeMO in cell-type-specific ATAC-seq peak regions. Lineage-specific motifs learned by the cell-type-specific ChromBPNet models are highlighted in red.
**(d)** Performance comparison of two LASSO regression models (9CT-All and 9CT-PeakOnly) predicting sequence conservation (PhyloP scores) from hematopoietic regulatory context features and local protein-coding gene constraints. Models were trained and evaluated with a leave-one-chromosome-out (LOCO) cross-validation scheme. Three evaluation sets are shown: genome-wide (all variants in the held-out chromosome), accessible (variants overlapping ATAC peaks in HSC/MPP/LMPP/CMP/CLP/pDC/GMP/MEP/mono), and inaccessible (variants outside peaks). Orange bars represent the 9CT-All model (29 features); blue bars represent the 9CT-PeakOnly model (11 features). Bars show the mean Pearson r across 22 chromosomes; error bars indicate ± s.e.m.; jittered points show individual chromosome correlations. Significance brackets denote paired t-tests across chromosomes, testing whether the 9CT-All model outperforms the 9CT-PeakOnly model (***P < 0.001, **P < 0.01, *P < 0.05, ns, not significant).
**(e-g)** S-LDSC enrichment of per-SNP heritability for blood cell traits in prioritized variants. Variants were ranked by predicted PhyloP conservation score and stratified into three thresholds: top 5%, top 1%, and top 0.1%. Two FLARE models are compared across facet panels (colors as indicated). Lines connect mean enrichment estimates across thresholds; shaded ribbons indicate 95% confidence intervals (±1.96 × s.e.m.). The red dashed horizontal line at 1 indicates the baseline (no enrichment). Pink dashed horizontal lines indicate the enrichment of the biologically expected progenitor cell type: MEP for trait RBC (e), CLP for trait LYM (f), and GMP for trait NEU (g).
**(h, i)** CADD score enrichment among variants prioritized by the two LASSO regression models (9CT-All and 9CT-PeakOnly). For each model, variants ranked by predicted FLARE score were stratified into top 0.1%, 1%, and 5%. Within each stratum, the odds ratio (OR) was computed using a one-sided Fisher's exact test (greater), comparing the proportion of variants exceeding CADD PHRED thresholds of 10 and 20 against the full background set. Horizontal bars represent the OR estimate; the dashed vertical line indicates OR = 1 (no enrichment). Asterisks denote Benjamini–Hochberg FDR-corrected significance (***P < 0.001, **P < 0.01, *P < 0.05), with exact FDR-corrected P values annotated to the right of each bar. A higher OR indicates that prioritized variants are more enriched for likely deleterious variants as scored by CADD.
